# A SREBF2-dependent gene program drives an immunotolerant dendritic cell population during cancer progression

**DOI:** 10.1101/2023.04.26.538456

**Authors:** Michael P. Plebanek, Yue Xue, Y-Van Nguyen, Nicholas C. DeVito, Xueying Wang, Alisha Holtzhausen, Georgia M. Beasley, Nagendra Yarla, Bala Thievanthiran, Brent A. Hanks

**Affiliations:** Department of Medicine Division of Medical Oncology Duke Cancer Institute Duke University Durham, NC 27708 USA; Department of Pharmacology and Cancer Biology Duke University Durham, NC 27708 USA; Lineberger Comprehensive Cancer Center University of North Carolina at Chapel Hill Chapel Hill, NC 27599 USA; Department of Surgery Division of Surgical Oncology Duke Cancer Institute Duke University Durham, NC 27708 USA

## Abstract

Dendritic cells (cDCs) are essential mediators of anti-tumor immunity. Cancers have developed mechanisms to render DCs dysfunctional within the tumor microenvironment. Utilizing CD63 as a unique surface marker, we demonstrate that mature regulatory DCs (mregDCs) suppress DC antigen cross-presentation while driving T_H_2 and regulatory T cell differentiation within tumor-draining lymph node tissues. Transcriptional and metabolic studies show that mregDC functionality is dependent upon the mevalonate biosynthetic pathway and the master transcription factor, SREBP2. Melanoma-derived lactate activates DC SREBP2 in the tumor microenvironment (TME) and drives mregDC development from conventional DCs. DC-specific genetic silencing and pharmacologic inhibition of SREBP2 promotes anti-tumor CD8^+^ T cell activation and suppresses melanoma progression. CD63^+^ mregDCs reside within the sentinel lymph nodes of melanoma patients. Collectively, this work describes a tumor-driven SREBP2-dependent program that promotes CD63^+^ mregDC development and function while serving as a promising therapeutic target for overcoming immune tolerance in the TME.

**One Sentence Summary:** The metabolic transcription factor, SREBF2, regulates the development and tolerogenic function of the mregDC population within the tumor microenvironment.

## INTRODUCTION

Dendritic cells (DCs) play a central role in orchestrating immune responses to cancers by presenting tumor antigens and promoting the activation of naïve T cells in tumor-draining lymph nodes (TDLNs) (*1, 2*). The ability of DCs to initiate adaptive responses is essential for the efficacy of immune checkpoint blockade (*3-5*), however, many cancers harbor dysfunctional DCs incapable of activating CD8^+^ T cells and propagating an immune response (*6, 7*). Indeed, multiple studies have demonstrated that tumor-associated DCs can exhibit a tolerogenic phenotype characterized by diminished antigen cross-presentation and an increased propensity to induce regulatory T cell (Treg) differentiation (*8-13*). Despite the well-established association of tolerogenic DCs and cancer, the detailed mechanisms that dictate their development and suppressive function have remained elusive.

Functional differences within the DC population have led to the classification of several distinct DC subsets, however a bona fide *in situ* tolerogenic DC subpopulation has not been definitively established. Conventional DCs (cDCs) arise from the common DC progenitor (CDP) and can further be divided into cDC1s recognized for their ability to cross-present antigen to stimulate CD8^+^ T cell responses and the more heterogenous cDC2s which are more proficient at activating various types of CD4^+^ T cell responses (*14*). More recently, the mature regulatory (mreg) cDCs which express high levels of DC maturation markers and immunoregulatory genes has been described (*15*). While mregDCs have been implicated in regulating tumor immunity, the roles of the mregDCs within the tumor microenvironment (TME) and their exact contributions to immune escape remain unclear (*15, 16*). Establishing the mechanisms leading to the development of mregDCs and how they influence the immune microenvironment has the potential to identify new drug targets and inform the design of novel immunotherapeutic strategies.

Recent studies have revealed that metabolic processes regulate the fundamental functional properties of DCs (*17, 18*). Upon DC activation, there is a shift in metabolism from fatty acid oxidation (FAO) and oxidative phosphorylation (OXPHOS) in resting DCs to aerobic glycolysis in order to rapidly generate ATP (*19*). This is coupled to increased fatty acid synthesis to support immunogenicity (*20*), but paradoxically, tumor infiltrating DCs with high lipid content exhibit impaired immunostimulatory properties (*7, 21-23*). In the context of peripheral tolerance, DC metabolism differs drastically and, like resting DCs, tolerogenic DCs utilize FAO for their energy demands (*24*). Critically, DC FAO promotes protoporphyrin IX synthesis and indoleamine-2,3-dioxygenase (IDO) activity which drives kynurenine production and Treg differentiation (*12*). Due to the importance of FAO, lipid homeostasis is tightly regulated by a complex signaling network. The sterol response element binding protein (SREBP) family of transcription factors are master regulators of lipid metabolism. In mammals, the SREBF family is composed of two genes, *SREBF1* and *SREBF2*. *SREBF*1 generates the isoforms SREBP1a and SREBP1c which are essential for fatty acid metabolism while *SREBF2* and its encoded protein SREBP2 controls cholesterol metabolism (*25*). This is accomplished by stimulating the expression of mevalonate (MVA) pathway genes including the target of statins, HMG-CoA reductase encoded by *HMGCR* to synthesize cholesterol, and the low-density lipoprotein (LDL) receptor-encoding gene, *LDLR,* which is responsible for the uptake of LDLs and extracellular cholesterol. Interestingly, the MVA pathway has previously been implicated in immunity (*26*). For example, patients with hyperimmunoglobin D syndrome (HIDS) have a loss of mevalonate kinase functionality and develop a pro-inflammatory condition (*27*). Additionally, studies have shown that inhibiting the MVA pathway promotes antigen presentation and enhances cytolytic T cell responses (*28, 29*). *SREBF2* expression and chromatin accessibility at SREBP transcription factor binding sites has been noted in cDC2s, but the importance of SREBF2 to DC biology in the context of disease remains unknown (*30*).

In the current study, we have identified a critical role for SREBP2 in regulating the development and function of mregDCs within the TME. These mregDCs can be isolated from other cDCs using the surface marker, CD63, and exhibit a reduced capacity to drive CD8^+^ T cell proliferation while enhancing Treg differentiation and suppressing T cell cross-priming by other DC subsets *in trans*. Further studies confirm that this tolerogenic CD63^+^ mregDC subset resides within the TDLN tissues of melanoma patients and maintains an enriched expression of genes involved in cholesterol homeostasis. This work further highlights the therapeutic targeting of SREBP2 and DC lipid metabolism as a promising approach to overcoming immune tolerance.

## RESULTS

### Identification of a tumor-associated DC population with a mevalonate pathway gene signature

DC activation and migration to the TDLN is necessary to support the generation of a robust anti-tumor immune response and the efficacy of checkpoint inhibitor immunotherapy (*31, 32*), however the mechanisms in which tolerogenic DCs regulate this process has remained unclear. To better understand the dynamic changes that occur in DCs within the TDLN during tumor progression, single-cell RNA sequencing (scRNA-seq) was applied to a BRAF^V600E^PTEN^-/-^ melanoma mouse model expressing a tamoxifen-inducible Cre recombinase driven by the tyrosinase promoter (*33*). Following the generation of a primary melanoma using this model, DCs (CD45^+^CD11c^+^MHCII^hi^F4/80^-^) from the inguinal TDLNs, distant non-draining lymph nodes (NDLNs), and inguinal LNs from non-tumor bearing control mice were enriched by FACS and analyzed by scRNA-seq to evaluate LN DC gene expression profiles (**Fig. S1A**). Utilizing Seurat v4 for graph-based unsupervised clustering, cDCs were identified based on the expression of *Flt3*, *Zbtb46*, and *Itgax* and further separated into cDC1 and cDC2 sub-populations by the expression of *Irf4*, *Irf8*, *Xcr1* and *Sirpa* (**Fig. 1A and Fig. S1B**)(*34*). During this analysis, a third cDC cluster was found to be enriched in TDLNs relative to NDLN and control LN tissues (**Fig. 1B, C and Fig. S1C**). Consistent with the previously reported mregDC subset, this cluster of DCs expresses elevated levels of immunosuppressive genes including *Cd274*, *Ido1*, *Ili41* and *Socs2* in addition to genes associated with DC maturation, such as *Cd40*, *Cd80*, *Cd83* and *Cd86* (**Fig. 1D and E**)(*15, 16, 35, 36*). Notably, however, this work found the mregDC subset to also exhibit a significant upregulation in genes associated with the mevalonate (MVA) biosynthetic pathway driven by the transcription factor SREBP2 (**Fig. 1F**). In support of this finding, gene set enrichment analysis (GSEA) identified that the hallmark cholesterol homeostasis gene set was highly enriched in mregDCs relative to other cDC populations (**Fig. 1G and Fig. S1D**). Importantly, these cDC gene expression profiles were found to be consistent between independent scRNAseq experimental replicates (**Fig. S1E**). We then utilized scATAC-seq to identify differentially accessible chromatin regions in these DC subpopulations (**Fig. 1H and Supplementary Data 1**). This analysis detected enriched accessible chromatin peaks in mregDCs that contain SREBP binding sites near several key genes, including *Hmgcr,* encoding the rate limiting enzyme of the MVA pathway, *Ldlr*, which encodes the low-density lipoprotein receptor, *Idi2,* which functions in the synthesis of cholesterol and isoprenoids, and *Fabp5*, a chaperone fatty acid binding protein (**Fig. 1I**). Overall, this work supports our previous finding of enhanced lipid metabolism in pro-tolerogenic DCs relative to other conventional DC subsets (*37*). Together, these data indicate that mregDCs accumulate within TDLN tissues during tumor progression and suggests that the MVA pathway and its master transcription factor, SREBP2, may play a role in regulating the development and function of this DC population.

**Fig. 1.**
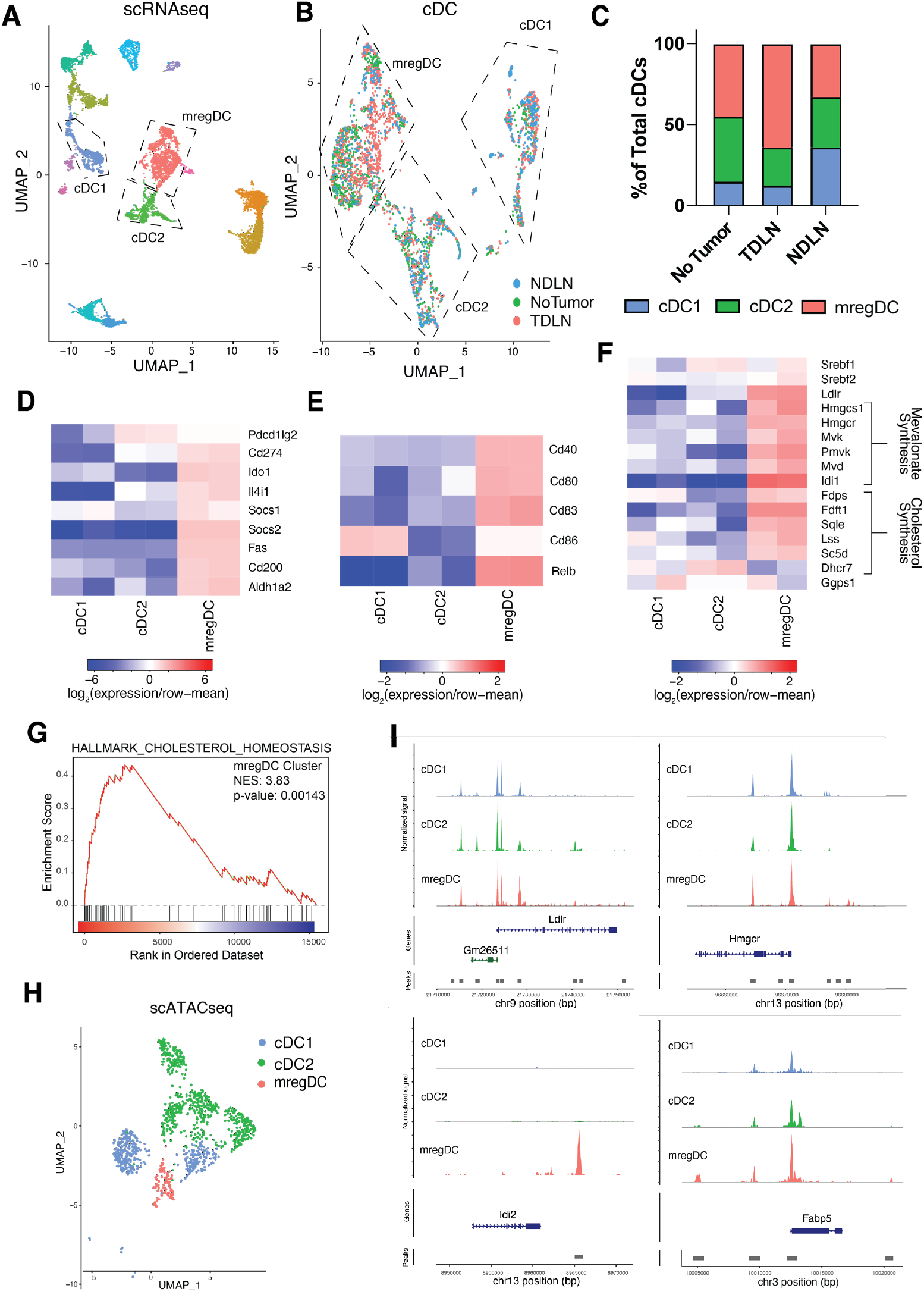
scRNA-seq identifies a conventional DC subset with increased expression of immunosuppressive and mevalonate pathway genes. (**A** to **G**) scRNA-seq performed on DCs sorted by FACS from the TDLNs and NDLNs of BRAF^V600E^ PTEN^-/-^ melanoma-bearing mice and the inguinal LNs of control non-tumor bearing mice (*n =* 2 independent experiments, ∼3000 cells per sample). (**A**) UMAP plot demonstrating clustering of DC sub-populations. (**B**) UMAP plot following re-clustering of cDCs showing the proportion of DCs in each sample. (**C**) cDC1, cDC2, and mregDC clusters as a percentage of cDCs in the TDLNs or NDLNs of melanoma-bearing mice and LNs of non-tumor bearing controls. (**D** to **F**) Heatmaps comparing cDC subset differential gene expression: (**D**) Key immunosuppressive genes, (**E**) DC maturation markers and (**F**) Mevalonate pathway genes. (**G**) GSEA of the Hallmark Cholesterol Homeostasis gene set in mregDCs relative to total DCs. (**H** and **I)** scATAC-seq performed on DCs sorted by FACS from the TDLNs of BRAF^V600E^ PTEN^-/-^ melanoma bearing mice (*n =* 2 independent experiments, ∼3000 cells per sample). (**H**) scATAC-seq UMAP plot demonstrating clustering of cDCs. (**I**) Chromatin accessibility tracks near the *Ldlr, Hmgcr, Idi2* and *Fabp5* genes in cDCs. All data is representative of 2-3 independent experiments.

### The tetraspanin CD63 serves as a marker of mregDCs which accumulate in TDLNs during tumor progression

To aid in the functional characterization of the mregDC sub-population, we proceeded to identify surface markers to distinguish mregDCs from other cDCs. Within the scRNAseq dataset, the gene *Cd63* encodes a surface tetraspanin that is highly enriched in the mregDC subset relative to other conventional DC populations (**Fig. 2A, B**). Next, we used flow cytometry to determine if increased *Cd63* gene expression by the mregDC subset corresponded with enhanced cell surface CD63 protein expression. Indeed, flow cytometry analysis demonstrates a CD63^hi^ population of DCs that can be effectively sorted from other conventional DC populations (**Fig. 2C and Fig. S2A**). Notably, relative to the CD63^-^ DC populations, CD63^+^ mregDCs express the cDC2 marker, CD172a, (SIRPα) to a greater degree than the cDC1 marker, XCR1. This is consistent with the scRNAseq data in which the mregDC subset expresses higher levels of the cDC2 genes, *Irf4* and *Sirpa,* relative to the cDC1 genes, *Irf8* and *Xcr1* (**Fig. 2C, D and Fig. S1B**). To confirm that the CD63^+^ DCs are representative of the mregDC sub-population, we used FACS to sort CD63^+^ DCs from the TDLNs of BRAF^V600E^PTEN^-/-^ melanoma-bearing mice and performed quantitative real-time polymerase chain reaction (qRT-PCR) analysis to quantify genes differentially expressed by mregDCs. This work confirmed elevated expression of *Cd63, Fabp5, Eno3* and *Socs2* by CD63^+^ DCs while the MVA pathway genes *Hmgcs1, Hmgcr, Pmvk, Mvd,* and *Idi1* were also significantly upregulated in CD63^+^ DCs and validated that SREBP2 gene targets are expressed at elevated levels in CD63^+^ mregDCs compared with CD63^-^ DCs (**Fig. 2E, F**). To further verify that CD63^+^ DCs were representative of the mregDC scRNAseq subset, we analyzed the expression of the DC chemokine receptor, CCR7, LDLR, and the maturation markers CD40, CD80 and CD86 by flow cytometry. Consistent with the scRNA-seq dataset (**Fig. 1**), CD63^+^ DCs express increased levels of CCR7, LDLR, CD40, CD80 and CD86 relative to other cDC subsets (**Fig. 2G**). This CD63^+^ mregDC population was further identified within the tumor tissues of the transgenic p53;Kras non-small cell lung cancer model indicating that this population is not unique to the BRAF^V600E^PTEN^-/-^ melanoma model (**Fig. S2B**).

**Fig. 2.**
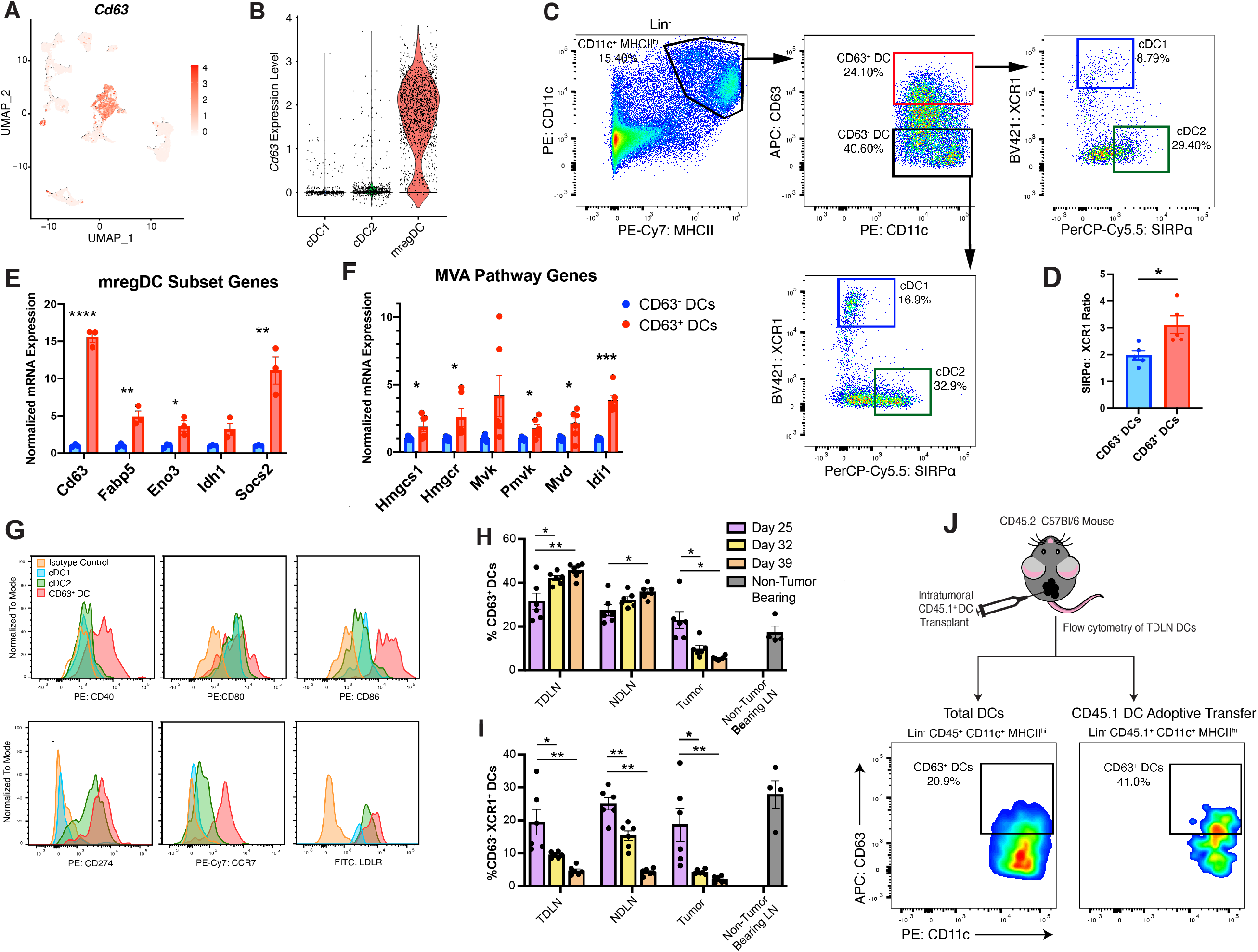
The tetraspanin CD63 is a marker of conventional mregDCs. (**A**) Tetraspanin *Cd63* gene expression overlaid on the UMAP plots generated in Fig. 1A (*n =* 2 independent experiments). (**B**) Violin plot showing gene expression of *Cd63* by cDC subset in the scRNA-seq dataset (*n =* 2 independent experiments). (**C**) Flow cytometry strategy to sort CD63^+^ DCs from LNs of tumor-bearing mice (*n =* 4). (**D**) Ratio of SIRPα^+^ cells to XCR1^+^ cells in CD63^-^ and CD63^+^ DC subsets (*n =* 4). (**E** and **F**) qPCR of FACS-sorted CD63^+^ or CD63^-^ DCs: (**E**) Key genes highly expressed in the mregDC scRNAseq cluster and (**F**) mevalonate pathway genes (*n =* 3). (**G**) Flow cytometry histograms demonstrating the expression of specific surface proteins from the mregDC scRNAseq cluster (*n =* 4). (**H** and **I**) Flow cytometry quantification of the percent of: (**H**) CD63^+^ DCs or (**I**) XCR1^+^ cDC1s in the TDLN, the NDLN, and the tumor of BRAF^V600E^ PTEN^-/-^ melanoma-bearing mice over time. Percentage of CD63^+^ DCs in the LNs of tumor naïve mice shown as a reference (*n =* 6). (**J**) Flow cytometry analysis of CD45.1^+^CD63^+^ DCs in the TDLN following intra-tumoral injection of isolated CD45.1^+^CD63^-^ DCs into CD45.2^+^ hosts harboring BRAF^V600E^PTEN^-/-^ melanomas (*n =* 4). All two-group comparisons were analyzed using unpaired *t* tests. Statistical analysis for three-group comparisons was performed using a two-way ANOVA followed by Sidak’s multiple comparisons test. All data presented as a mean value ± standard error of mean (SEM) and is representative of 2 independent experiments. \**P* < 0.05, \*\**P* < 0.005. \*\*\**P* < 0.0005.

Leveraging the CD63 surface marker, we utilized flow cytometry to investigate the dynamics of the CD63^+^ DC subset throughout tumor progression. Towards this end, the CD63^+^ DC subset was quantified in tumor, TDLN, and NDLN tissues at multiple time points following tumor induction in the transgenic BRAF^V600E^ PTEN^-/-^ mouse model. In support of the previous scRNAseq data, flow cytometry demonstrates an increase in CD63^+^ DCs in the TDLN relative to the NDLN and the LN of non-tumor bearing mice (**Fig. 2H)**. Importantly, this work further showed the increase in the CD63^+^ DC population within TDLNs to coincide with a reduction of this population within the tumor bed during the course of tumor progression. Consistent with a potential immunosuppressive role for mregDCs in tumor immunity, we further found increased numbers of CD63^+^ DCs to correlate with a relative decrease in the antigen cross-presenting cDC1 subset within TDLN tissues (**Fig. 2I**). An increased proportion of CD63^+^ DCs also reside within LNs relative to the primary tumor which is consistent with the migratory phenotype of mregDCs (**Fig. S2C**). Also, the gene expression profiles were noted to be stable when comparing cDC populations in TDLNs to DCs from LNs of non-tumor bearing mice, suggesting that the mregDC subset is likely not developing in the LN but trafficking from other tissues (**Fig. S2D**).To test the hypothesis that CD63^+^ DCs are generated in the tumor bed prior to trafficking to LN tissues, CD45.1^+^CD63^-^ DCs were delivered into the tumor bed of BRAF^V600E^PTEN^-/-^ melanoma-bearing CD45.2^+^ hosts. Flow cytometry analysis revealed a significant number of CD45.1^+^CD63^+^ DCs within TDLNs, confirming that tumor infiltrating DCs can undergo conversion into CD63^+^ DCs in the tumor bed prior to migrating to TDLN tissues (**Fig. 2J and Fig. S2E**). Together, these data suggests that CD63^+^ DCs develop in the tumor bed and migrate to TDLN tissues during tumor progression.

### CD63^+^ mregDCs suppress CD8^+^ T cell responses and promote Treg differentiation

After finding that CD63^+^ mregDCs accumulate within TLDN tissues, we were interested in understanding the role of these DCs in directing adaptive immunity. Towards this goal, we were initially interested in quantifying the ability of tumor-derived CD63^+^ DCs to drive CD8^+^ T cell proliferation relative to CD63^-^ DCs. We therefore co-cultured sorted CD63^+^ or CD63^-^ DCs from the TDLNs of mice bearing implanted syngeneic BRAF^V600E^ PTEN^-/-^ tumors engineered to express ovalbumin (OVA) with naïve CD8^+^ T cells isolated from OT-I T cell receptor (TCR) transgenic mice. This work demonstrated CD63^+^ DCs to exhibit a diminished capacity to stimulate antigen-specific CD8^+^ T cell responses relative to CD63^-^ DCs (**Fig. 3A**). Consistent with these findings, we also used a flow cytometry-based conjugated OVA assay to measure impaired antigen uptake and processing by CD63^+^ DCs relative to cDC2s subsets (**Fig. S3A**). This observed impairment in T cell activation was likely due to defects in antigen cross-presentation pathways, as the CD63^+^ DCs maintained their capacity to directly present the SIINFEKL peptide and stimulate OT-I T cell responses (**Fig. 3B**).

**Fig. 3.**
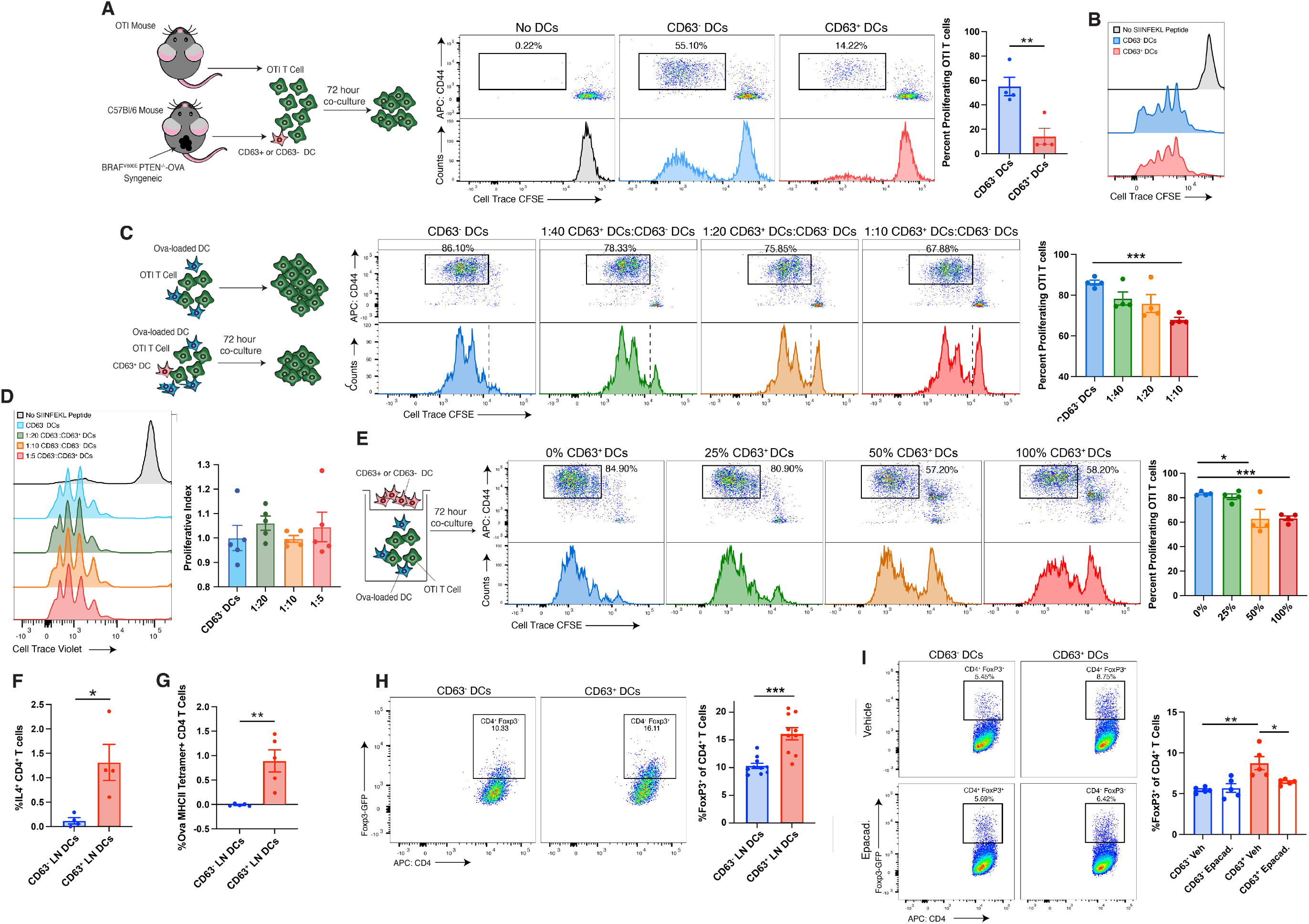
CD63^+^ DCs suppress CD8^+^ T cells and promote CD4^+^ Treg differentiation. (**A** to **E**) T cell proliferation assays with CFSE-labeled OTI T cells co-cultured with antigen-pulsed DCs. (**A**) Co-culture of OTI CD8^+^ T cells with either CD63^-^ or CD63^+^ DCs isolated from the TDLN of mice bearing OVA-expressing BRAF^V600E^PTEN^-/-^ melanomas (5:1 T cells: DC) (Left) Schematic of experiment (Right) Flow cytometry plots and quantification (*n =* 4). (**B**) Co-culture of OTI CD8^+^ T cells with either CD63^-^ or CD63^+^ DCs pulsed with SIINFEKL peptide prior to co-culture. (**C**) Co-culture of OTI CD8^+^ T cells with OVA-pulsed DCs and increasing numbers of CD63^+^ DCs (Left) Schematic of experiment (Right) Flow cytometry plots and quantification (*n =* 4). (**D**) Co-culture of OTI CD8^+^ T cells with SIINFEKL-pulsed DCs and increasing numbers of CD63^+^ DCs (*n =* 5). (**E**) Co-culture of OTI CD8^+^ T cells with SIINFEKL-pulsed DCs (1 x 10^4^) in the bottom chamber of a 0.4 μm transwell. Ratios of CD63^+^:CD63^-^ DCs ranging from 1 x 10^4^:0 to 0:1 x 10^4^ in upper chamber while maintaining the same total cell number across conditions. (**F**) Intracellular staining of IL-4 in CD4^+^ T cells following co-culture with CD63^+^ or CD63^-^ DCs (T cell:DC ratio, 5:1) (*n =* 4). (**G**) MHCII-OVA_323-329_ tetramer staining of CD4^+^ T cells following co-culture with CD63^+^ or CD63^-^ DCs isolated from OVA-expressing melanoma-bearing mice (T cell:DC ratio, 5:1) (*n =* 5). (**H**) Flow cytometry plots (left) and quantification (right) of Foxp3-expressing Tregs after co-culture of CD63^-^ or CD63^+^ DCs (H-2^d^) with naïve CD4^+^ T cells isolated from Foxp3-GFP mice (H-2^b^) (T cell:DC ratio, 5:1) (*n =* 10). (**I**) Flow cytometry plots (left) and quantification (right) demonstrating the effect of the IDO1 inhibitor, epacadostat, on CD63^+^ DC-induced Treg differentiation (*n =* 5). All two-group comparisons were analyzed using unpaired *t* tests. (**C** to **E** and **I**) Statistical analysis was performed by two-way ANOVA followed by Sidak’s multiple comparisons test. All data presented as a mean ± SEM. All data is representative of 2-3 independent experiments. \**P* < 0.05, \*\**P* < 0.005. \*\*\**P* < 0.0005.

Rather than simply being less efficient in antigen processing and CD8^+^ T cell activation, we investigated whether CD63^+^ DCs actively suppress the generation of effector T cell responses. We performed additional CD8^+^ T cell response assays utilizing CD63^+^ DCs co-cultured at various ratios with OVA-pulsed CD63^-^ DCs. These studies demonstrated that at even lower ratios of CD63^+^ DCs:CD63^-^ cDCs (1:10), antigen cross-presentation by nearby cDCs can be effectively suppressed by CD63^+^ DCs *in trans* (**Fig. 3C**). This suggests that a relatively small number of CD63^+^ DCs can potently impair local CD8^+^ T cell activation. Also, it is notable that this inhibitory effect targets DC antigen cross-presentation as the CD63^+^ DCs are unable to suppress the direct antigen presentation of the SIINFEKL peptide by nearby cDCs *in trans* (**Fig. 3D**). We then performed CD63^+^ DC:CD63^-^ DC co-culture assays using a 0.4 μm transwell system that tests the impact of CD63^+^ DC-derived soluble molecules on DC antigen cross-presentation *in trans*. This work also demonstrated inhibition of OT-I CD8^+^ T cell responses, confirming that CD63^+^ DCs inhibit antigen cross-presentation by nearby DCs via the production of a soluble mediator (**Fig. 3E**).

Interestingly, while CD63^+^ DCs exhibited a limited ability to promote CD8^+^ T cell proliferation, CD63^+^ DCs sorted from BRAF^V600E^PTEN^-/-^ tumor-bearing mice maintained a robust capacity to promote OT-II CD4^+^ T cell proliferation (**Fig. S3B**). Additional flow cytometry studies revealed these expanded CD4^+^ T cells to exhibit a T_H_2 phenotype (**Fig. 3F**). This finding is consistent with increased expression of T_H_2-associated genes by the mregDCs cluster in the scRNAseq dataset (**Fig. S3C**). Additionally, MHC class II tetramer co-culture studies utilizing CD63^+^ mregDCs isolated from mice harboring OVA-expressing tumors, demonstrated CD63^+^ mregDCs to expand tumor antigen-specific CD4^+^ T cells more readily than CD63^-^ control DCs (**Fig. 3G**). This data raised the possibility that the CD63^+^ mregDC population may support the differentiation of FoxP3^+^ Tregs. It is well established that the expression of tryptophan degrading enzymes such as IL4I1 and IDO1 by myeloid cells within the TME can promote the differentiation of Tregs (*38-40*) and, the expression of these enzymes was found to be enriched in the CD63^+^ DC population relative to cDC1 and cDC2s (**Fig. 1D**). Consistent with these findings, co-cultures of allogeneic naïve CD4^+^ T cells with CD63^+^ DCs reveal an enhanced capacity to promote CD4^+^FoxP3^+^ Treg differentiation relative to CD63^-^ DCs in an IDO1-dependent manner (**Fig. 3H and I**). Collectively, these data suggest that CD63^+^ mregDCs accumulate within TDLN tissues and promote the generation of an immunotolerant microenvironment.

### CD63^+^ mregDCs can be resolved into two distinct sub-populations

Since our findings suggest that mregDCs exhibit a pro-tolerogenic function, we were interested in understanding the developmental relationship between mregDCs and other cDC subsets. We, therefore, re-clustered the cDC populations and generated cDC signature scores by selecting the top differentially expressed genes by each DC subtype within our scRNAseq dataset. Upon higher resolution analysis, the mregDCs separated into two sub-populations which we refer to as mregDC2 and mregDC1 based on their transcriptional signature scores (**Fig. 4A and B**). Notably, the mregDC2 cluster was highlighted by the expression of the lipid chaperone, *Fabp5* and the oxysterol receptor *Gpr183*, and was found to be more closely associated with the cDC2 genetic program while the mregDC1 subset was determined to be more closely associated with the cDC1 subset (**Fig. 4C, D and Supplementary Table 2**). To examine the origins of the mregDC populations, we performed pseudotime analysis to determine the developmental trajectory between the conventional DC subpopulations. Indeed, consistent with the mregDC gene expression profile, this approach suggested that there is a developmental trajectory originating in the cDC2 cluster that branches into both mregDC subsets (**Fig. 4E**). To confirm these findings, we isolated cDC2s and cDC1s from the spleens of CD45.1 expressing mice by FACS and delivered these cells into the BRAF^V600E^PTEN^-/-^ tumors of CD45.2^+^ hosts prior to harvesting TDLN tissues for further analysis. This work confirmed that CD45.1^+^ cDC2s differentiate more readily into the CD45.1^+^CD63^hi^ mregDC subset within the TME compared with CD45.1^+^ cDC1s (**Fig. 4F**). Overall, these findings indicate that the mregDC population is phenotypically heterogenous and develops predominantly from cDC2s.

**Fig. 4.**
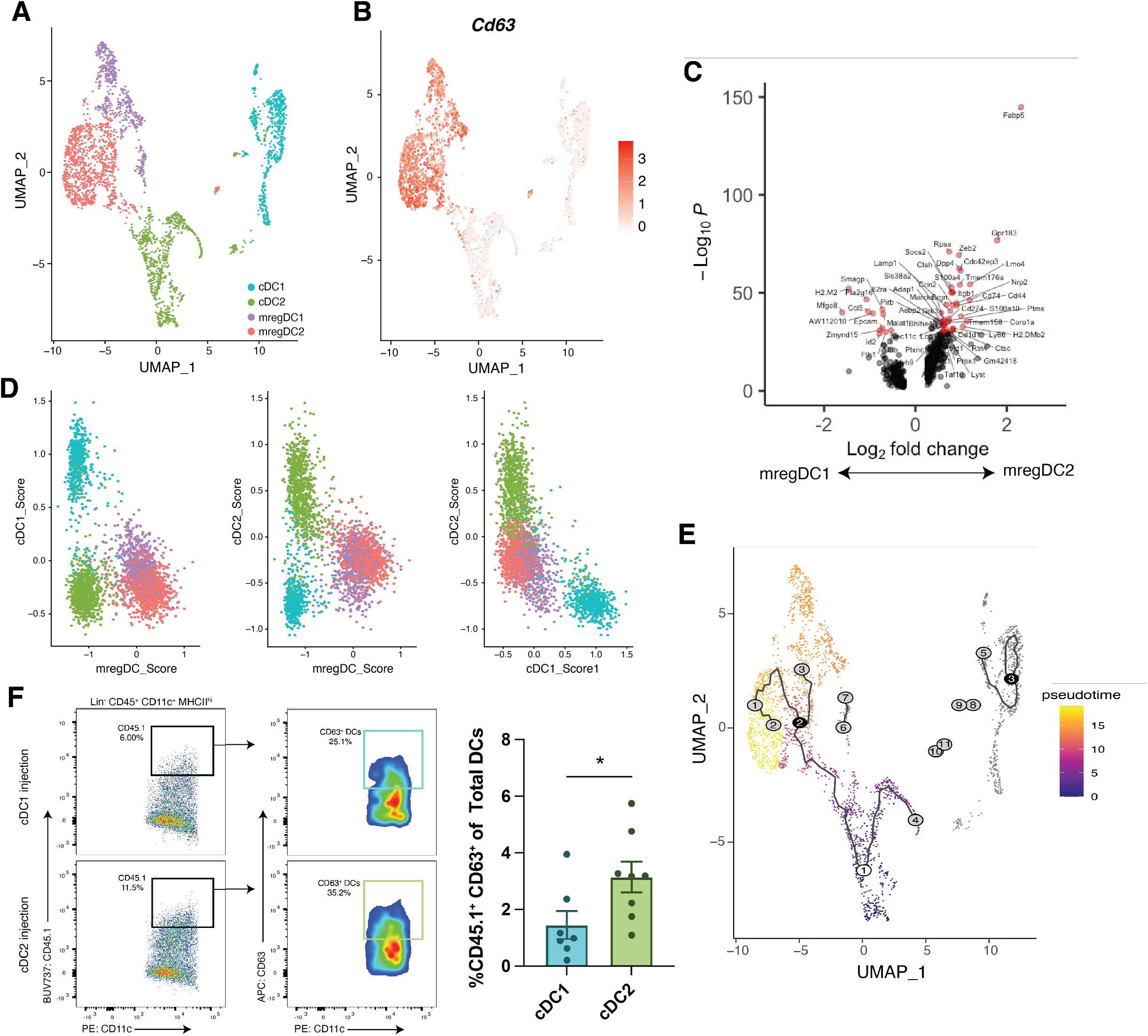
CD63^+^ mregDCs express a distinct genetic signature and can be divided into two sub-populations. (**A**) UMAP plot showing a subset of the scRNAseq data from **Fig. 1** displaying only the conventional DC clusters. (**B**) Gene expression overlaid on UMAP plots showing *Cd63* expression in cDC clusters from **A**. (**C**) Volcano plot displaying the differentially expressed genes between the mregDC1 and mregDC2 subpopulations. (**D**) DC subset scores were generated from the top differentially expressed genes in the original clusters from Fig. 1 and applied to the different DC sub-populations comparing the mregDC subsets to cDC1 and cDC2. (**E**) Pseudo-time analysis displaying the cellular trajectory of cDC2s toward mregDCs. (**F**) (Left) Flow cytometry plots of CD45.1^+^ CD63^+^ DCs following intra-tumoral injection of (top) CD45.1^+^ cDC1s or (bottom) CD45.1^+^ cDC2s into CD45.2^+^ hosts. (Right) Quantification of the percent CD45.1^+^CD63^+^ DCs of the total DCs following intra-tumoral injection of CD45.1^+^ cDC1 or cDC2 (*n =* 8). (**F**) Data analyzed using unpaired *t* test. All data presented as a mean ± SEM. All data is representative of 2 independent experiments. \**P* < 0.05, \*\**P* < 0.005, \*\*\**P* < 0.0005.

### mregDCs are metabolically distinct from cDCs and exhibit elevated levels of FAO

As described in prior studies, metabolism is a critical determinant of DC function (*18*), and mregDCs are enriched in the expression of multiple enzymes in the MVA metabolic pathway driven by the transcription factor, SREBP2 (**Fig. 1F**). It has been demonstrated that tumor-infiltrating DCs have increased lipid body formation and increased lipid peroxidation which can promote abnormal lipid synthetic programs and ultimately interfere with DC antigen cross-presentation (*7, 21, 41*). As a result, we proceeded to analyze and compare the metabolic profiles of the CD63^+^ mregDC subset with other cDC populations. First, we tested whether these DC sub-populations contained different amounts of neutral and oxidized lipids. Relative to cDC1s and cDC2s, we found that CD63^+^ mregDCs harbored an increase in both neutral and oxidized lipid content as indicated by incorporation of BODIPY 493/503 and oxidized BODIPY 581/591, respectively (**Fig. 5A and B)**. Altered neutral lipid content could be supported by increased lipid uptake from the surrounding TME or through increased flux through the MVA pathway leading to the synthesis of cholesterol. The transcription factor, SREBP2, is known to play a role in each of these processes by promoting expression of the MVA pathway genes and the *LDLR* gene responsible for cholesterol uptake (*42, 43*).

**Fig. 5.**
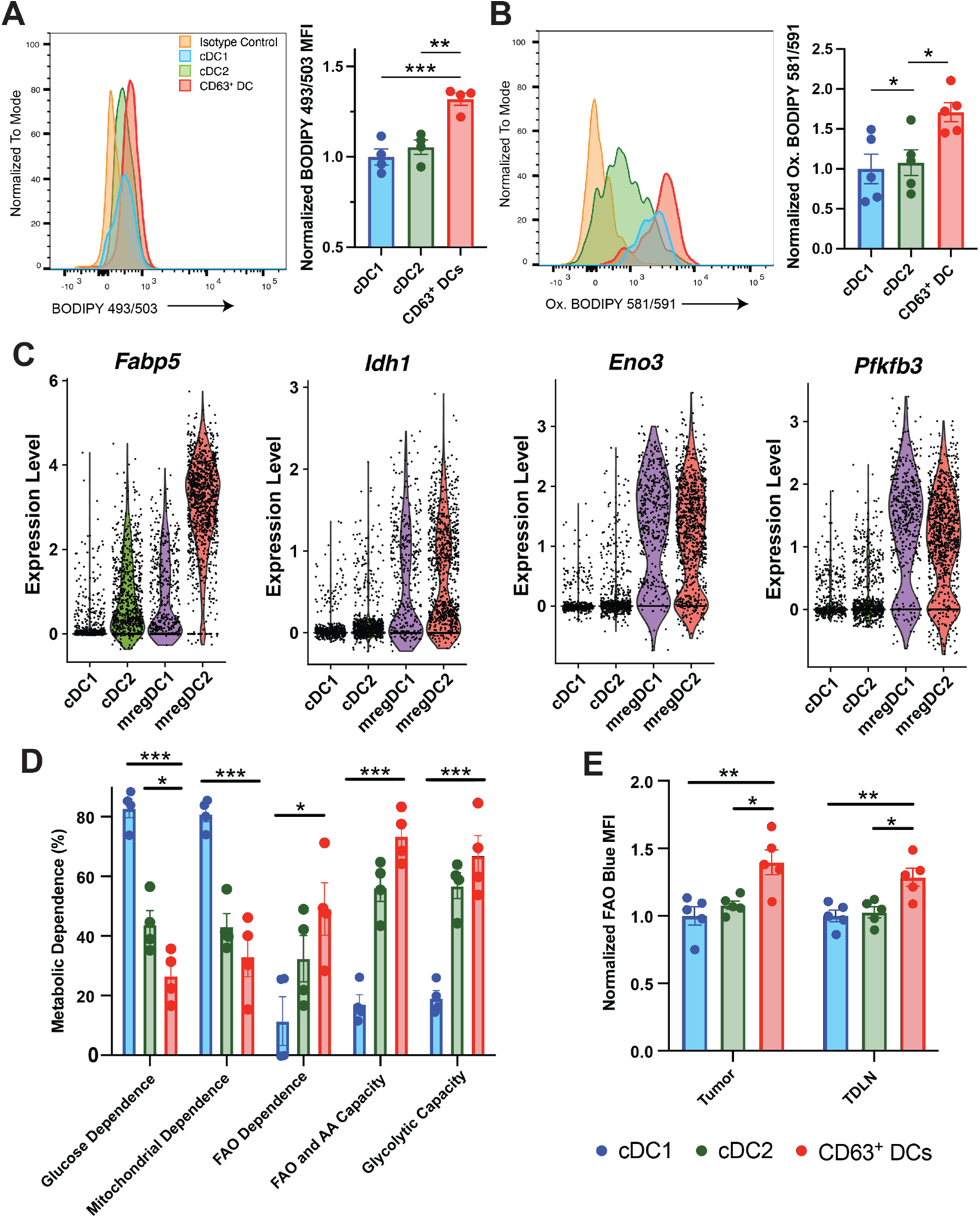
CD63^+^ mregDCs are metabolically distinct. (**A**) Flow cytometry analysis of cDC neutral lipid content based on BODIPY 493/503 flow cytometry. (Right) Representative flow cytometry histogram of BODIPY 493/503 mean fluorescence intensity (MFI) (*n =* 4). (**B**) Flow cytometry analysis of cDCs oxidized lipid content based on BODIPY 581/591 C11 flow cytometry. (Right) Representative flow cytometry histogram of BODIPY 581/591 C11 MFI (*n =* 5). (**C**) Transcriptional analysis of critical metabolic genes in the cDC subpopulations derived from the scRNA-seq analysis in **Fig. 1** (*n =* 2 independent experiments). (**D**) SCENITH assay analysis of cDC subpopulations (*n =* 4). (**E**) Flow cytometry quantification of FAOblue staining in cDCs. Data normalized to cDC1 FAOblue MFI (*n =* 5). Statistical analysis was performed by two-way ANOVA followed by Sidak’s multiple comparisons test. Data presented as a mean ± SEM. All data is representative of at least two independent experiments. \**P* < 0.05, \*\**P* < 0.005. \*\*\**P* < 0.0005.

The MVA pathway functions to convert acetyl-CoA generated from glycolysis or FAO into cholesterol and isoprenoids (*44*). Due to the increased expression of MVA pathway enzymes by CD63^+^ mregDCs (**Fig. 1F**) and alterations in mregDC lipid content, we further generated metabolic pathway gene expression scores using the KEGG database and compared central metabolic pathway gene expression in mregDCs to cDC1s and cDC2s. This analysis found mregDCs to exhibit relatively elevated expression of several rate limiting enzymes, including *Idh1* which catalyzes the rate-limiting step of the TCA cycle, *Eno3* and *Pfkb3,* two rate limiting glycolytic enzymes, as well as *Fabp5*, which encodes a fatty acid chaperone (**Fig. 5C and Fig. S4A-D**). While probing for gene expression is informative, it does not provide functional information regarding the metabolic pathways that are important regulators of these DC subpopulations. Therefore, to determine the predominant metabolic pathways employed by mregDCs, we used the flow cytometry-based SCENITH method which measures protein synthesis/puromycin incorporation as a surrogate for ATP generation (*45*). This approach showed CD63^+^ mregDCs were more dependent on FAO and exhibit an increased capacity for FAO/amino acid oxidation relative to other cDC populations (**Fig. 5D and Fig. S5A, B**). Following this analysis, it was still unclear if CD63^+^ mregDCs were actively metabolizing fatty acids to a greater degree than other DC subsets. To test this, we used a fatty acid derivative (FAOblue) that fluoresces following three cycles of the mitochondrial FAO system. This assay demonstrated an increase in FAO in CD63^+^ mregDCs in the tumor and TDLN relative to both cDC1 and cDC2 subsets (**Fig. 5E**). All together, these experiments indicate that mregDCs are metabolically distinct from other cDC populations and exhibit enhanced levels of FAO. These findings are consistent with our previous work linking elevated levels of FAO with DC-mediated Treg differentiation and the induction of immune tolerance in melanoma (*37*). Based on these findings, we initiated a series of studies to explore the impact of CD63^+^ mregDCs on tumorigenesis.

### SREBP inhibition reverses CD63^+^ mregDC tolerance and suppresses tumor progression

SREBP2 is a critical transcription factor that regulates the expression of MVA pathway genes (*46, 47*). Since lipid homeostasis plays such a critical role in regulating the functionality of CD63^+^ mregDCs, we hypothesized that SREBP2 inhibition would interfere with the ability of CD63^+^ mregDCs to regulate T responses. To initially test this hypothesis, CD63^+^ or CD63^-^ DCs were isolated from mice and treated with the SREBP inhibitor, fatostatin. Co-cultures of fatostatin-treated CD63^+^ DCs with allogeneic naïve CD4^+^ T cells exhibited a reduced capacity to promote Treg differentiation relative to untreated CD63^+^ DCs (**Fig. 6A**). Additionally, fatostatin as well as the HMG-CoA reductase inhibitor, simvastatin, induced a significant increase in OT-I T cell cross-priming by CD63^+^ mregDCs relative to vehicle treated controls (**Fig. 6B**). Interestingly, the geranyl-geranyl transferase inhibitor, GGTI-2147, was unable to increase cross-priming, suggesting that the downstream isoprenoid biosynthesis pathway does not regulate CD63^+^ mregDC activation of effector T cells (**Fig. 6B**). Together, these results led us to hypothesize that SREBP inhibition would reverse CD63^+^ mregDC immunosuppression and promote an anti-tumor immune response *in vivo*. To test this hypothesis, autochthonous BRAF^V600E^PTEN^-/-^ melanomas were treated with fatostatin (30mg/kg) twice weekly once the tumors reached approximately 75 mm^3^. While fatostatin treatment significantly slowed the proliferation of BRAF^V600E^PTEN^-/-^ melanoma cells *in vitro,* it also reduced expression of SREBP2 target genes in CD63^+^ mregDCs sorted from fatostatin treated mice (**Fig. S6A and B**). Importantly, SREBP2 inhibition with fatostatin led to a significant reduction in tumor growth compared to vehicle treated controls (**Fig. 6C**). Accompanying flow cytometry studies reveal that fatostatin diminishes CD63^+^ mregDCs while increasing cDC1s numbers in both the TDLN and tumor tissues (**Fig. 6D**). These findings correlated with a suppression in Treg differentiation and a reciprocal increase in the numbers of CD8^+^ T cells within TDLN and tumor tissues (**Fig. 6E and Fig. S6C, D**). Although our studies confirm that fatostatin exhibits tumor-intrinsic inhibitory properties, these findings further indicate that systemic SREBP2 inhibition generates a TME that supports enhanced anti-tumor immunity. Next, to verify the importance of DC SREBP2 for tumor progression, we utilized CD11c promoter (*Itgax)*-driven Cre recombinase mice to generate a DC-specific SREBF2 knock-out mouse model (**Fig. S7A and B**). Consistent with its role in mitochondrial-dependent respiration, BMDCs from these CD11c-SREBF2^-/-^ mice also demonstrate a diminished oxygen consumption rate as shown by extracellular flux analysis and decreased puromycin incorporation based on SCENITH assays, correlating with lower ATP production (**Fig. 6F and Fig. S7C**). In support of SREBP2s role in regulating lipid homeostasis, the neutral lipid content of BMDCs differentiated from CD11c-SREBF2^-/-^ mice was significantly less than the BMDCs derived from control SREBF2^fl/fl^ littermates (**Fig. S7D)**. Similarly, BMDCs differentiated from CD11c-SREBF2^-/-^ mice demonstrated diminished FAO relative to control mice (**Fig. 6G)**. We performed additional assays to determine the influence of SREBP2 on DC-dependent antigen cross-presentation. BMDCs from CD11c-SREBF2^-/-^ or control SREBF2^fl/fl^ littermates were differentiated and loaded with OVA prior to co-culture with naive CD8^+^ T cells isolated from OTI mice, demonstrating that SREBP2^-/-^ DCs have an increased capacity to promote CD8^+^ T cell proliferation (**Fig. 6H).** These findings suggest that SREBF2^-/-^ DCs exhibit enhanced T cell stimulation capacity and could therefore suppress tumor progression. To test this hypothesis, we implanted syngeneic BRAF^V600E^PTEN^-/-^ melanomas into CD11c-SREBF2^-/-^ and control SREBF2^fl/fl^ littermates and monitored tumor growth for 25 days. This study demonstrated suppressed tumor growth as well as diminished numbers of CD63^+^ mregDCs and a concurrent increase in cDC1s in the TME of CD11c-SREBF2^-/-^ mice relative to SREBF2^fl/fl^ control mice (**Fig 6I and J**). These alterations were further found to be associated with a reduction in tumor-infiltrating CD4^+^ T_H_ cells and Tregs as well as an increase in CD8^+^ T cells within the TME (**Fig 6K**). Additionally, hematoxylin and eosin staining as well as immunohistochemical analysis of the melanoma-associated antigen, S100β, of resected lung tissues was performed to assess for evidence of metastatic progression to the lung. These studies demonstrated fewer pulmonary micro-metastases in CD11c-SREBF2^-/-^ hosts compared to control SREBF2^fl/fl^ littermates (**Fig. 6L**). Overall, these findings provide further support that SREBP2 serves as a key regulator of mregDC-mediated immune suppression while contributing to tumor progression.

**Fig. 6.**
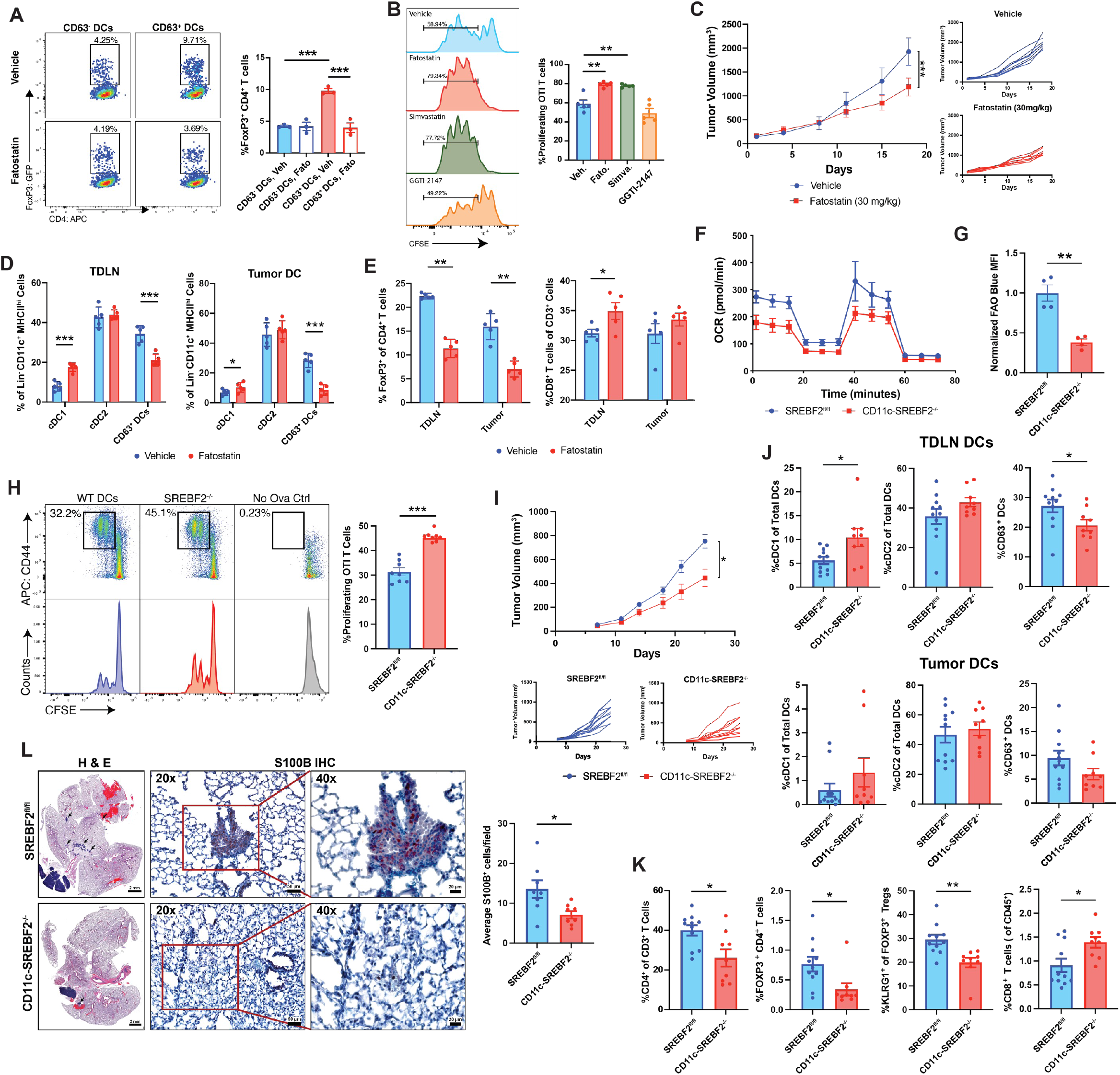
SREBP2 is a critical transcription factor for CD63^+^ mregDCs. (**A**) Flow cytometry analysis following CD63^+^ or CD63^-^ DC co-culture with FoxP3-GFP naïve CD4^+^ T cells pre-treated with Fatostatin or vehicle control. (Left) Flow cytometry dot plots (Right) Quantification of flow cytometry plots (*n =* 3). (**B**) Flow cytometry CFSE proliferation assay following OVA-pulsed CD63^+^ or CD63^-^ DCs co-cultured with naïve OTI CD8^+^ T cells ± fatostatin, simvastatin, or GGTI-2147. (Left) Flow cytometry histograms. (Right) Quantification of flow cytometry plots (*n =* 5). (**C**) (Left) Tumor growth curves of primary melanoma tissues in autochthonous BRAF^V600E^PTEN^-/-^ transgenic melanoma mice treated with fatostatin (30mg/kg) or vehicle control. (Right) Growth curves showing individual replicates (*n =* 9). (**D** to **E**) Flow cytometry quantification of immune cell populations within TDLN or tumor tissues following fatostatin versus vehicle control treatment of an autochthonous BRAF^V600E^PTEN^-/-^ transgenic melanoma model. (**D**) Quantification of DC subtypes in the (Left) TDLN and (Right) Tumor. (**E**) Quantification of (Left) Foxp3^+^ Tregs and (Right) CD8^+^ T cells in the TDLN and Tumor. (**F**) Extracellular flux analysis measuring oxygen consumption rate (OCR) in CD11c-SREBF2^-/-^ control littermate SREBF2^fl/fl^ hosts. (*n =* 5). (**G**) Flow cytometry quantification of FAOblue staining in BMDCs isolated from CD11c*-Cre*, *Srebf2^-/-^* mice compared to littermates lacking Cre expression (SREBF2^fl/fl^). Data normalized to cDC1 FAOblue MFI (*n =* 4). (**H**) Flow cytometry CFSE proliferation assay following OVA-pulsed BMDCs from CD11c-SREBF2^-/-^ mice and their SREBF2^fl/fl^ littermate controls co-cultured with naïve OTI CD8^+^ T cells. (Right) Quantification of flow cytometry plots (*n =* 8). (**I**) (Top) Tumor growth curves of BRAF^V600E^PTEN^-/-^ melanomas in CD11c-SREBF2^-/-^ mice and their SREBF2^fl/fl^ littermate controls. (Bottom) Growth curves showing individual replicates (*n =* 12). (**J** and **K**) Flow cytometry quantification of tumor and TDLN infiltrating immune cell populations in CD11c-SREBF2^-/-^ mice and their SREBF2^fl/fl^ littermate controls. (**J**) Quantification of DC subtypes in the (Above) TDLN and (Below) Tumor. (**K**) Quantification of tumor-infiltrating CD4^+^ T cells, Foxp3^+^ CD4^+^ Tregs, KLRG1^+^-activated Tregs and CD8^+^ T cells (*n =* 11 for SREBF2^fl/fl^ controls, *n =* 9 for CD11c-SREBF2^fl/fl^). (**l**) CD11c-SREBF2^-/-^ mice and their SREBF2^fl/fl^ littermate controls were implanted with YUMM1.1g BRAF^V600E^PTEN^-/-^CDKN2A^-/-^ melanoma cells and allowed to grow for 28 Days. Lungs were sectioned and H&E and S100β IHC was used to quantify micro-metastases. All two-group comparisons were analyzed using unpaired *t* tests. (**A** and **B**) Statistical analysis was performed by two-way ANOVA followed by Sidak’s multiple comparisons test. Data is shown as mean ± SEM with individual data points. All data is representative of 2-3 independent experiments.* *P* < 0.05, \*\**P* < 0.005, \*\*\**P* < 0.0005.

### Tumor-derived lactate promotes SREBP2 activation in the tumor microenvironment

In prior experiments, we have demonstrated that mregDCs are enriched in neutral lipids relative to cDC1s and cDC2s (**Figs 5A,B**). Conventionally, under high sterol states, SREBP2 is tethered to the ER preventing it from activating its target genes(*48*). Despite this, an SREBP2 target gene signature is enriched in mregDCs relative to cDC1s and cDC2s, indicating that there are alternative mechanisms of SREBP2 activation. An acidic pH (pH 6.8) within the tumor microenvironment can promote SREBP2 activation, and nuclear translocation, leading to the expression of SREBP2 target genes involved in cholesterol homeostasis (*49*). Melanomas have a notably high rate of aerobic glycolysis leading to the acidification of the TME via the production of lactic acid (*50, 51*). Therefore, we reasoned that melanoma-produced lactate could promote SREBP2 activation in DCs. To test this hypothesis, BMDCs were treated with lactate (50 mM, pH 6.8) and the nuclear translocation of SREBP2 was measured demonstrating increased SREBP2 activation (**Fig. 7A**). Next, BMDCs from either CD11c-SREBF2^-/-^ mice or control SREBF2^fl/fl^ littermates were treated with lactate (50 mM, pH 6.8) or left untreated (pH 7.4) and RT-qPCR was performed. Lactate treatment led to increased expression of SREBP2 target genes including *Ldlr, Hmgcs1, Hmgcr,* and *Idi1* (**Fig. 7B**). Importantly, the increased expression of these genes was prevented in DCs lacking SREBP2 expression. These experiments demonstrate that lactate can promote the expression of genes involved in cholesterol homeostasis and that this result is dependent upon SREBP2. Based on these results, we were interested in exploring whether tumor-derived lactate within the TME could promote the development of CD63^+^ mregDCs. We therefore used CRISPR-Cas9 to delete the lactate exporter MCT1 (*Slc16a1)* from a BRAF^V600E^PTEN^-/-^ melanoma cell line **(Fig. S7E**). Lactate secretion was found to be significantly diminished in the conditioned media harvested from BRAF^V600E^PTEN^-/-^Slc16a1^-/-^ cells **(Fig. S7F**). As a result, BRAF^V600E^PTEN^-/-^Slc16a1^-/-^ melanoma cells were implanted in syngeneic mice and tumor growth as well as CD63^+^ mregDC development within TDLN tissues was monitored. BRAF^V600E^PTEN^-/-^Slc16a1^-/-^ melanoma growth was significantly diminished relative to wild-type control tumors (**Fig. 7C**). Additionally, flow cytometry also demonstrated that mice bearing BRAF^V600E^ PTEN^-/-^ Slc16a1^-/-^ melanomas were associated with fewer CD63^+^ mregDCs within the TDLNs than mice bearing BRAF^V600E^PTEN^-/-^ control melanomas (**Fig. 7D**). These data indicate that tumor-derived lactate leads to SREBP2 activation in DCs within the TME and that this mechanism drives the development of tolerogenic CD63^+^ mregDCs.

**Fig. 7.**
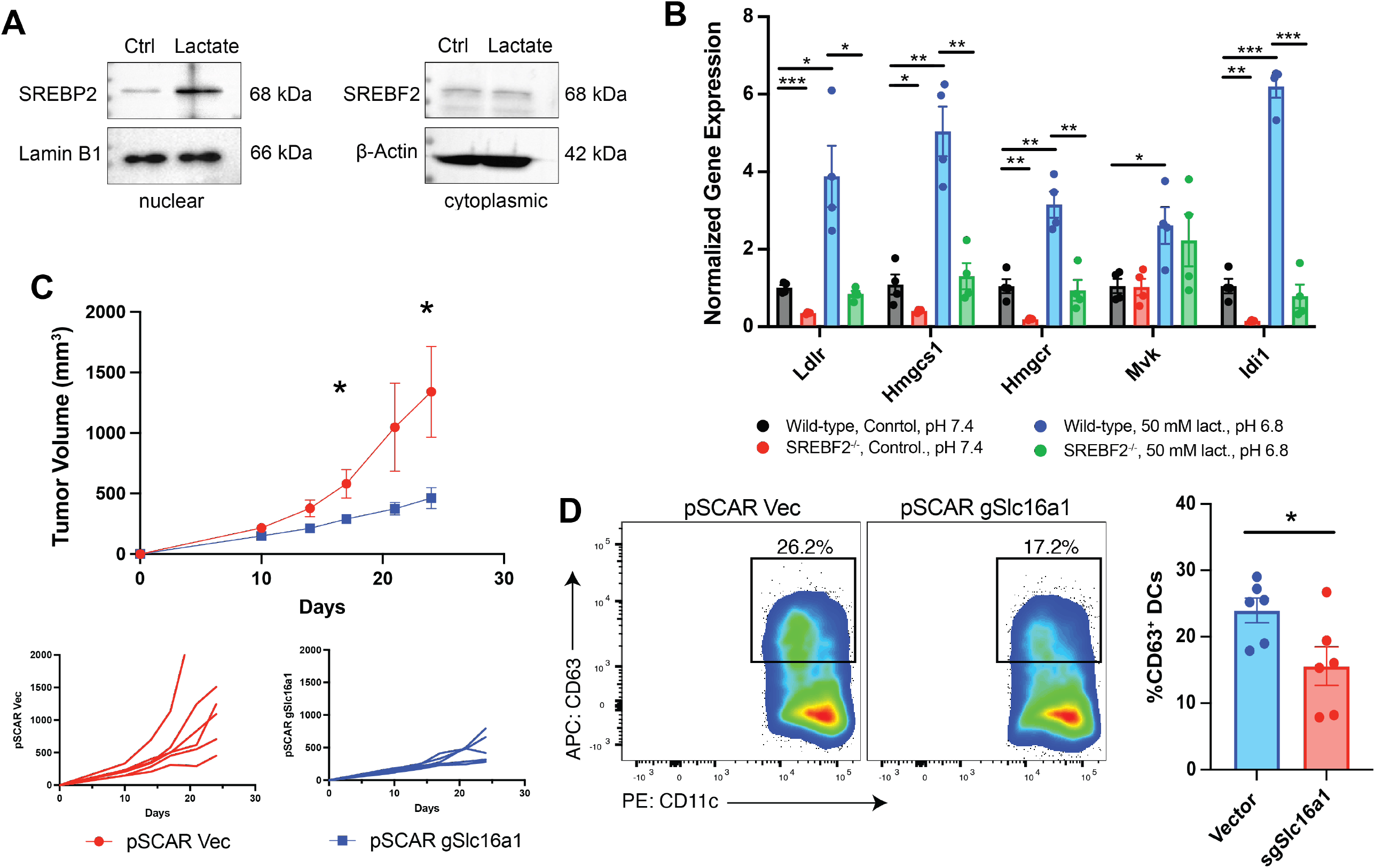
Tumor-derived lactate promotes SREBP2 activation in the tumor microenvironment. (**A**) Western blot showing activation and nuclear translocation of SREBP2 following treatment of DCs with lactate 50 mM (pH 6.8). (**B**) qRT-PCR of BMDCs isolated from CD11c-SREBF2^-/-^ mice or their SREBF2^fl/fl^ littermate controls following treatment with lactate (50 mM. pH 6.8) or under normal conditions (pH 7.4) probing for the SREBP2 target genes *Ldlr, Hmgcs1 Hmgcr, Mvk,* and *Idi1* (n=4). (**C**) (Top) Tumor growth curves of BRAF^V600E^PTEN^-/-^ melanomas with CRISPR-Cas9 mediated knockout of *Slc16a1* or non-edited controls. (Bottom) Growth curves showing individual tumor progression (*n =* 6). (**D**) Flow cytometry quantification of CD63^+^ DCs infiltrating the TDLN in mice bearing *Slc16a1^-/-^* or non-edited control BRAF^V600E^PTEN^-/-^ melanoma cells (n=6). All two-group comparisons were analyzed using unpaired *t* tests. (**B**) Statistical analysis performed by two-way ANOVA followed by Sidak’s multiple comparisons test. Data is shown as mean ± SEM with individual data points. All data is representative of 2-3 independent experiments. * *P* < 0.05, \*\**P* < 0.005, \*\*\**P* < 0.0005.

### CD63^+^ mregDCs are conserved in the sentinel LNs of melanoma patients

Given our pre-clinical data suggesting that the CD63^+^ mregDC population plays a critical role in supporting the development of an immunotolerant TME, we investigated human sentinel LN tissues harvested from melanoma patients undergoing sentinel LN biopsy procedures for the presence of this DC population. We initially performed immunofluorescence (IF) microscopy which suggested the presence of CD63^+^CD11c^+^ DCs in harvested human LN tissue specimens (**Fig. 8A**). Using the CD63^+^ DC expression score that we previously established, we also collected fresh human sentinel LN tissue and utilized scRNAseq to confirm the presence of this CD63^+^ mregDC subset (**Fig. 8B and C**). Following confirmation that mregDCs existed in the sentinel LNs of melanoma patients, we proceeded to determine if these mregDCs were enriched in the cholesterol homeostasis gene expression similar to their mouse counterparts. Indeed, GSEA demonstrates increased enrichment of the Hallmark Cholesterol Homeostasis gene set relative to the other cDC clusters (**Fig. 8D**). In order to assess the spatial location of the CD63^+^ mregDC population within the tissue architecture of human LN tissues relative to Treg and CD4^+^ T cell populations, we performed spatial transcriptional analysis which confirmed the presence of the CD63^+^ mregDC subset within the paracortex of human LN tissues (**Fig. 8E**). Notably, the mregDC expression score positively correlated with CD4^+^ T cell and CD4^+^ Treg expression scores indicating mregDC co-localization with these T cell populations within human LN tissues (**Fig. S8A and B**).

**Fig. 8.**
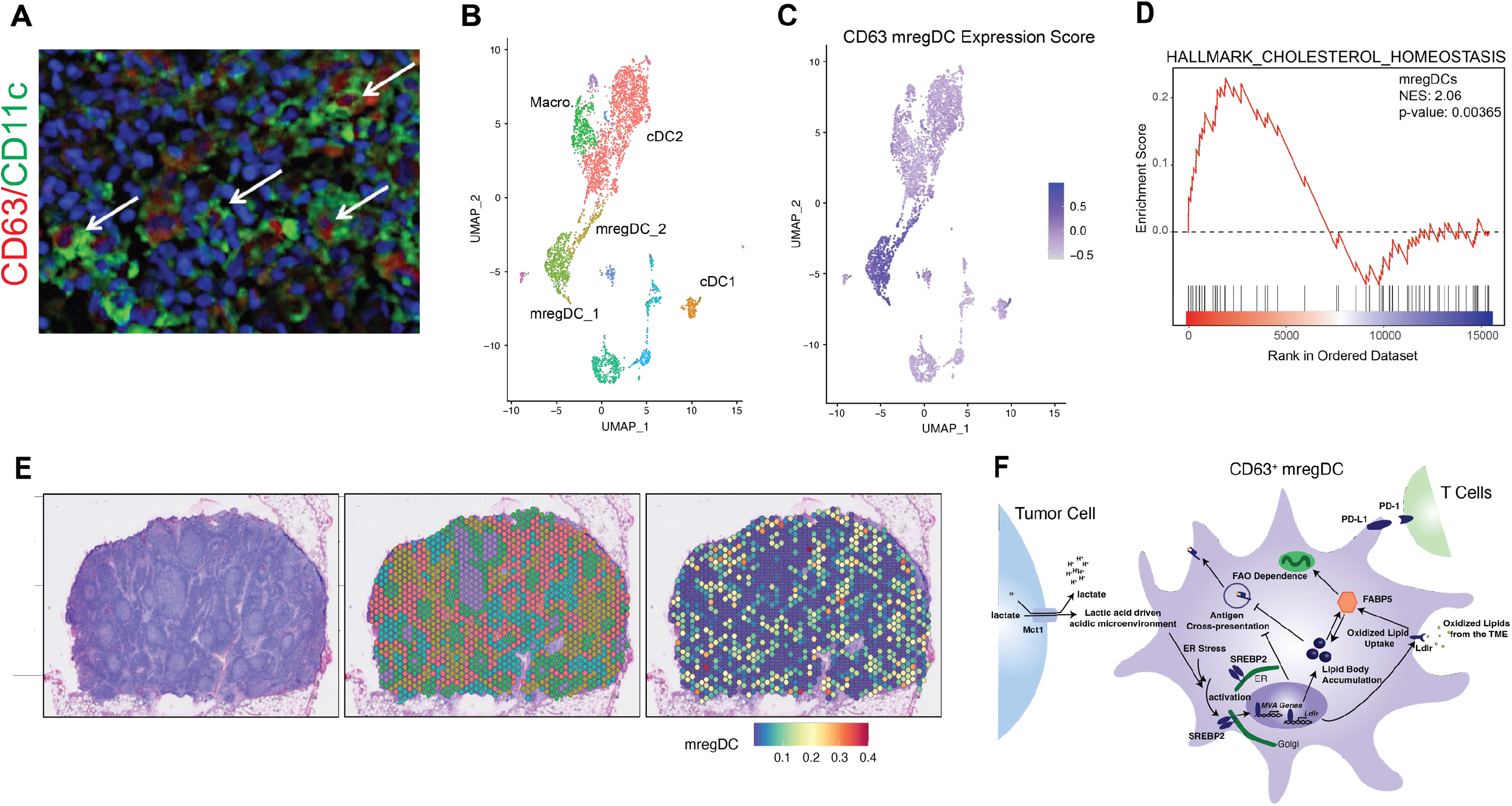
CD63^+^ DCs are conserved in humans. (**A**) Immunofluorescence staining of CD63 (red) and CD11c (green) sentinel LN sections isolated from melanoma patients. (*n =* 3 independent experiments). White arrows, CD63^+^CD11c^+^ cells. (**B**) scRNA-seq UMAP plot demonstrating clustering of DC sub-populations sorted from the sentinel LN of a melanoma patient. (*n =* 2 independent experiments). (**C**) CD63^+^ mregDCs expression score generated in **Fig. 4** applied to the human melanoma scRNA-seq UMAP plot. (**D**) Gene set enrichment analysis (GSEA) of the Hallmark Cholesterol Homeostasis gene set in human mregDCs relative to total DCs. (**E** and **F)** Visium spatial transcriptomic analysis of sentinel LN tissues harvested from melanoma patients. (**E**) (left) H&E staining of sentinel LN tissue. (middle) Visium spots overlaid on H&E image. (right) mregDC expression score overlaid on the Visium spatial transcriptomics spot map demonstrating the presence of mregDCs in human melanoma patients (*n =* 3 independent experiments). (**G**) Schematic showing the role of SREBP2 in regulating mregDCs. All data is representative of 2-3 independent experiments.

## DISCUSSION

Despite recent advances in oncology, a significant percentage of cancer patients remain refractory to checkpoint inhibitor immunotherapy (*52, 53*). There have been various tumor-dependent characteristics reported to influence immunotherapy responses (*54, 55*). However, our understanding of tumor-mediated immune evasion mechanisms remains incomplete and our ability to overcome anti-PD-1 immunotherapy resistance has remained limited. Although DCs are recognized as being a key mediator of anti-tumor immunity by facilitating the activation of effector CD8^+^ T cells, there has been a growing appreciation that evolving tumors can co-opt select DC populations to generate a tolerant tumor microenvironment (*9, 56, 57*). We propose that an improved understanding of tolerogenic DCs will serve as a foundation for the development of a novel family of therapeutic agents capable of expanding immunotherapy responsive cancer patient populations.

A distinct DC subset with inherent tolerogenic properties within the TME termed ‘mature DCs enriched in immunoregulatory molecules’ (mregDCs) was recently reported and shown to drive T_H_2 polarized immune responses capable of suppressing anti-tumor immunity in pre-clinical tumor models(*15*). We now expand on this report by characterizing the development of mregDCs within the TME, the metabolic pathways that regulate mregDC function, and the underlying mechanisms that enable mregDC populations to suppress anti-tumor immunity. We demonstrate that the mevalonate biosynthetic pathway and its master transcription factor, SREBP2, plays a critical role in supporting the development and function of the CD63^+^ mregDCs (**Fig. 8F**). By identifying CD63 as a restrictive surface marker of the mregDC subset, we further show that mregDCs possess a potent immunoregulatory function allowing them to drive Treg development while also suppressing DC antigen cross-presentation *in trans*, even at high conventional DC:CD63^+^ mregDC ratios. Finally, we demonstrate that both pharmacologic SREBP2 inhibition and DC-specific *Srebf2* silencing reverses mregDC-dependent immune evasion while also suppressing tumor progression *in vivo*. IF microscopy, scRNAseq, and spatial transcriptional analysis was further utilized to demonstrate that the CD63^+^ mregDC population resides within the draining LN tissues of melanoma patients. Together, these findings support a role for SREBP2-dependent mregDCs in establishing an immunotolerant tumor microenvironment and suggests that this pathway represents a promising target for suppressing tumor progression and overcoming immunotherapy resistance.

In light of the important role of metabolism in regulating DC function, we initiated studies to determine the unique metabolic characteristics of CD63^+^ mregDCs relative to other DC subsets (*24, 58*). Our findings indicate that CD63^+^ mregDCs uniquely exhibit enhanced levels of expression of several MVA enzymes as well as their master transcription factor, SREBP2. Additional studies have further revealed CD63^+^ mregDCs to harbor increased lipid stores as well as enhanced levels of FAO relative to both type 1 and 2 cDCs. The CD63^+^ mregDC subset also exhibits elevated expression of the LDL receptor (LDLR) and fatty acid binding protein-5 (FABP5), previously shown to suppress IL-12 expression (*59*). These results are consistent with previous studies linking the MVA synthesis pathway with inflammatory regulation (*60, 61*). This is highlighted by the reported clinical autoinflammatory syndromes associated with MVK deficiency (*62, 63*). Prior work has explained this relationship by suggesting that impaired MVA flux induces the stimulation of STING-mediated type I interferon signaling as well as the activation of the pyrin inflammasome (*64, 65*). Still, a more recent study has determined that the inhibition of the MVA pathway using either statins or bisphosphonates can promote antigen presentation via increased antigen processing and endosomal recycling (*66*). Indeed, this is consistent with our findings showing that both SREBP2 and HMG-CoA reductase inhibition enhanced CD63^+^ mregDC-mediated activation of CD8^+^ T cell responses. Additional retrospective studies have also demonstrated statin use to be associated with improved clinical outcomes for cancer patients undergoing anti-PD-1 immunotherapy (*67, 68*). Our results suggest that the pharmacologic inhibition of CD63^+^ mregDC function likely contributes to these observed clinical outcomes.

Importantly, this work also suggests that these metabolic pathways have critical functional implications for CD63^+^ mregDCs as both SREBP2 inhibition with fatostatin and DC-specific genetic silencing of *Srebf2* enhances CD8^+^ T cell activation, reduces Treg differentiation, and suppresses *in vivo* BRAF^V600E^PTEN^-/-^ melanoma progression. As the geranylgeranyltransferase-I inhibitor of downstream isoprenoid biosynthesis, GGTI-2147, failed to effectively enhance CD63^+^ mregDC-mediated CD8^+^ T cell responses, our studies suggest that the MVA pathway contributes to these immunologic effects via the production of cholesterol. Regardless, these findings suggest that SREBF2 is a key central regulator of the immunosuppressive properties of CD63^+^ mregDCs. Indeed, prior studies have shown SREBP2 to be induced by the PI3K-Akt pathway, the same pathway known to be involved in vitamin D3-mediated induction of DC tolerance (*69-71*). Based on these cumulative studies, further investigation into the role of the PI3K-Akt pathway in regulating SREBP2 function in CD63^+^ mregDCs is warranted. Overall, these results are consistent with our prior data, as well as the data of others, implicating FAO as a critical component of the molecular machinery necessary for tumor-mediated DC tolerization (*37, 72*).

While this work contributes to an improved understanding of the underlying biology of tolerogenic DCs, additional work will be necessary to fully elucidate the pathways involved in SREBP2-mediated regulation of the downstream transcriptional program of CD63^+^ mregDCs. While conditional SREBF2 deletion demonstrated the importance of SREBP2 to mregDC development, the conditional deletion utilized a *ITGAX* promoter-driven Cre recombinase resulted in unrestricted loss of SREBF2 across all DC subtypes including mregDCs. Notably, we were unable to use a ZBTB46 driven Cre as SREBF2 loss in these mice was surprisingly embryonically lethal. In view of the role of SREBP1 in lipogenesis, additional studies are also necessary to differentiate its role from SREBP2 in directing CD63^+^ mregDC function.

The underlying mechanisms regulating the development of the mregDC subset within the TME also remains unclear. Our data indicates that CD63^+^ mregDCs expand and migrates from the tumor bed to local draining LN tissues throughout the course of tumor progression. Our data further shows that CD63^+^ mregDCs serve to potently impair effector CD8^+^ T cell responses while promoting both Treg and T_H_2 CD4^+^ T cell differentiation, creating a tolerized immune microenvironment. While prior authors have described mregDCs as differentiating both from type 1 and type 2 cDC populations, our studies including transcriptional profiling, pseudo-time trajectory analysis, and adoptive transfer experiments indicate that a greater number of mregDCs derive from type 2 cDCs (*15, 73*). We conjecture that this finding may explain the functional disparity reported for the mregDC subset, describing both a CD8^+^ T cell activation role and a capacity to support pro-tumorigenic T_H_2 polarized immune responses (*15, 74*). Despite this insight, it was still uncertain exactly how tumors induce the differentiation and expansion of the CD63^+^ mregDC population within the TME. Based on prior studies implicating a role for low pH in the activation of SREBP2, we engineered BRAF^V600E^PTEN^-/-^ melanomas genetically silenced for the primary lactate transporter, MCT1, and examined its impact on the regulation of anti-tumor immunity and tumor progression. Indeed, these studies confirmed that tumor-dependent lactate production within the TME plays an important role in driving the development of CD63^+^ mregDCs via the activation of SREBP2-dependent signaling.

Previous work has highlighted the important role that tolerogenic DCs play in creating immunosuppressive microenvironments that are conducive for tumor growth. Studies illustrating the potency of these tolerogenic DCs in the inhibition of anti-tumor immunity have shown that only small numbers of tolerogenic DCs are necessary for suppressing effector T cell responses (*10, 75, 76*). Our findings indicate that the inhibition of antigen cross-presentation by nearby DCs via the release of a soluble mediator may explain this phenomenon. Considering the critical role that DC-dependent antigen cross-presentation plays in the efficacy of anti-PD-1 immunotherapy, we propose that this mechanism may represent a key target for enhancing immunotherapy responses in traditionally refractory tumors (*3, 5, 77*). Additional studies will be necessary to identify the CD63^+^ mregDC-derived soluble factor(s) mediating the suppression of antigen cross-presentation by cDC1s.

Herein, we provide new insight into the biochemical pathways underpinning the function of the mregDC population while highlighting their dynamics and supportive role in tumor progression. Overall, this work provides a foundation for future studies investigating and leveraging the mregDCs for augmenting anti-tumor immunity and overcoming resistance to available checkpoint inhibitor immunotherapies.

## Materials and Methods

### Experimental Mice

Mice were housed and maintained in an isolated animal facility by the Duke University’s Division of Laboratory Animal Resources (DLAR) in accordance with IACUC approved animal protocols. C57Bl/6 mice (Strain #000664), C57Bl/6-Tg(TcraTcrb)1100Mjb/J (OT-1, Strain #003831), B6.Cg-Foxp3^tm2Tch^/J (Foxp3-GFP, Strain #006772) and BALB/c mice (Strain #000651) B6.Cg-Tg(Tyr-cre/ERT2)13Bos BRAF^tm1Mmcm^ Pten^tm1Hwu^/BosJ (BRAF^CA^;PTEN^lox/lox^; Tyr::CreERT2, Strain #013590) were purchased from The Jackson Laboratory and bred in our animal colony. To generate dendritic cell specific *Srebf2* KO mice, B6.Cg-Tg(Itgax-cre)1-1Reiz/J (Strain #008068, CD11c-Cre) mice were crossed to Srebf2^tm1.1Jdh^/J (Strain #031792, Srebf2^fl^). All experiments randomly assigned 6- to 8-week-old littermates and both male and female mice were included for all experiments.

### Cell Culture

The BRAF^V600E^ PTEN^-/-^ melanoma cell line was generated by spontaneous immortalization of a tumor resected from the BRAF^CA^;PTEN^lox/lox^;Tyr::CreERT2 mouse model (*13*) and cultured in DMEM containing 10% FBS (Genesee Scientific, Cat# 25-550) and 1% antibiotic-antimycotic (ThermoFisher Cat. #15240062). The cytotoxic effect of fatostatin on the BRAF^V600E^ PTEN^-/-^ melanoma cell line was evaluated using a colorimetric MTS assay. As per the manufacturer’s instructions (Promega, Cat# G3582) the absorbance at 490 nm was determined 1 hour following the addition of MTS and its inhibitory concentration 50 (IC_50_) was determined at 24 and 48 hours after treatment with fatostatin.

Bone marrow dendritic cells were generated by flushing the tibia and fibula of C57Bl/6 mice with RPMI according to a previously established protocol (*78*). Bone marrow cells were then separated from stroma, red blood cells and debris by centrifuging at 800 x g on top of Lympholyte Cell Separation Media (Cedarlane Labs, CL5030) and cultured in RPMI containing 10% FBS, 1% anti-anti, 2-mercaptoethanol, 20 ng/ml GM-CSF (Peprotech, 315-03) and 10 ng/ml IL-4 (BioAbChem, 42-IL4 C). Media was replenished every 48 hours for 7 days at which point BMDCs were harvested and magnetically selected using CD11c specific microbeads (Miltenyi Biotec, Cat# 130-125-835).

### qRT-PCR

RNA was isolated from DCs using RNeasy Micro kit (Qiagen, Cat# 74004) and cDNA was synthesized using iScript cDNA Synthesis Kit (Biorad, Cat# 1708890). For low input reactions, cDNA libraries were pre-amplified using Sso Advanced PreAmp Supermix (Biorad, Cat#1725160). qRT-PCR was performed using Power SYBR Green PCR Master Mix (ThermoFisher Scientific, Cat# 4367659) on an Applied Biosystems AB7500 real time PCR instrument. The delta-delta Ct method was used to analyze and report qRT-PCR results.

### Western Blot

Protein lysates were extracted using RIPA Lysis and Extraction Buffer (ThermoFisher, 89901) supplemented with complete protease inhibitor and phosphatase inhibitor (Roche, 4693159001 and 4906845001). Protein samples were separated by SDS-PAGE and transferred onto PVDF membranes (Bio-Rad). Primary antibodies and appropriate horseradish peroxidase (HRP)- conjugated secondary antibodies (Jackson Immuno Research Laboratories) were used for blotting. The proteins were visualized by ECL-Plus (GEHealthcare) using Image Quant LAS500 (GE Healthcare Life Sciences). The following antibodies were used: anti-SREBP2 (Novus Biologicals, NB100-74543) 1:1000 overnight at 4°C, anti-MCT1 (Novus Biologicals, 20139-1-AP) 1:1000 overnight at 4°C, anti-β-Actin (C4, Santa Cruz, sc-47778) 1:1000 for 1 hour at room temperature).

### Flow Cytometry

For flow cytometry of tumor infiltrating immune cells, tumors were resected and single cell suspensions were generated by incubating in a 5 mL solution of RPMI containing collagenase type IV (1 mg/mL, Sigma Aldrich Cat. C5138), Hyaluronidase Type V (0.1 mg/mL, Sigma Aldrich Cat. H6254) and DNase Type IV (20 U/mL), Sigma Aldrich Cat. D5025) for 30 min. at 37° C with mechanical dissociation every 10 minutes using a gentleMACs Tissue Dissociator (Miltenyi Biotec). Following the incubation, the digested tumor samples were strained through 40 μm cell strainers and the digestion quenched with 10 mL of RPMI containing 10% FBS. The cells were then washed in PBS and the RBCs were lysed using RBC lysis buffer (5 min, room temperature). Following RBC lysis cells were washed in PBS and then stained for FACS.

For flow cytometry of lymph node cells, LNs were resected and digested in a solution of a 5 mL solution of RPMI containing collagenase type IV (1 mg/mL, Sigma Aldrich Cat. C5138) and DNase Type IV (20 U/mL) Sigma Aldrich Cat. D5025) for 20 minutes at 37°C. Cells were then strained through 40 μm cell strainers and the digestion quenched with 10 mL of RPMI containing 10% FBS. Cells were then washed in PBS and resuspended to a dilution of 5 x 10^6^ cells/mL for FACS staining.

For FACS staining, first cells were stained with Live/Dead Fixable Aqua or Violet Dead Cell Stain Kit. (Thermo Fisher Scientific, L34957/L34955) at a concentration of 1 μl Live Dead stain/ml for 30 minutes in PBS at room temperature. Following the incubation, cells were washed 2x in FACS buffer (5% FBS, 2 mM EDTA in PBS) and resuspended with anti-CD16/CD32 Mouse BD FC Block (BD Biosciences, Cat. 553142) in FACS buffer at a concentration of 10 ug/ml for 15 min. at 4°C. Following FC block, fluorophore conjugated primary antibodies were added to cells in FACS buffer and incubated for 1 hour at 4°C. Cells were washed 2x in FACS buffer and fixed using BD Cytofix (BD Biosciences, Cat. 554655) for 15 min at 4°C then washed an additional 2 times and resuspended in FACS buffer. Cells were analyzed on a BD Fortessa X-20 flow cytometer at the Duke Cancer Institute Flow Cytometry Core Facility.

### Immunohistochemistry and Immunofluorescence Analysis

Paraffin sections (5 μm) from lung tissues were processed using standard protocols for immunohistochemistry (IHC) staining according to manufacturer’s instructions (Abcam: ab236468). Tissues were permeabilized by incubation in 0.4% Triton-X in TBS for 20 min, hydrogen peroxide block for 10 min and protein block for 1 hour. Anti-S100β (rabbit polyclonal antibody, 1:500, Novus biological, NBP1-87102) was incubated at 4^0^C overnight followed by application of a rabbit-specific HRP/3-amino-9-ethylcarbazole (AEC) IHC Detection Kit - Micro-polymer (abcam, ab236468) incubated at room temperature for 1 hr. Sections were imaged with an Axio Imager upright microscope. Area calculations for S100β-staining of metastatic foci were quantified at 40X magnification and averaged over 3 sections per specimen. Dual immunofluorescence (IF) was performed on paraffin-embedded tissues that were sectioned at 5 μm. Tissue sections were labeled for the following antigens using a dual IF assay: CD63 (NBP2-32830, Novus Biologicals) and CD11c (PA0554, Leica). This reaction was carried out on the Bond Rx fully automated slide staining system (Leica Biosystems) using the Bond Research Detection kit (DS9455). Slides were dewaxed in Bond Dewax solution (AR9222) and hydrated in Bond Wash solution (AR9590). Heat induced antigen retrieval was performed at 100°C in Bond-Epitope Retrieval solution 1 pH 6.0 (AR9961) for either 20 or 10 minutes. After pretreatment, tissues were blocked, and primary antibodies were diluted as follows: CD63 at 1:500 and CD11c at 1:2. Ready-to use secondary antibodies Novolink Post Primary and Novolink Polymer (RE7260-CE, Leica) were used followed by either TSA Cy5 (SAT705A001EA, Akoya Biosciences) or TSA Cy3 (SAT704A001EA, Akoya Biosciences) to visualize the target of interest. Nuclei were stained with Hoechst 33258 (Invitrogen). The stained slides were mounted with ProLong Gold antifade reagent (P36930, Invitrogen).

### scRNA-seq Processing and Analysis

For scRNA-seq experiments, 5,000 DCs were sorted from the tumor draining lymph node (inguinal) and the distant lymph node (axillary) of BRAF^V600E^ PTEN^-/-^ melanoma bearing mice or from the inguinal lymph node of non-tumor bearing mice using FACS as previously described. Single cells were encapsulated in gel bead-in-emulsions (GEMs) using the Chromium Next GEM Single Cell 3’ kit v3.1 (10x Genomics, Cat. PN-1000269) according to the manufacturer’s instructions. Quality control of libraries was performed using an Agilent 2100 Bioanalyzer and the final NGS libraries were quantified using NEBNext Library Quant Kit for Illumina (New England Biolabs, Cat. E7630L). For NGS, samples were sequenced on an Illumina NovaSeq 6000 at a depth of ∼40,000 reads per cell.

For the scRNA-seq analysis, sequencing reads were demultiplexed and processed using the Cell Ranger 4.0.0 (10x Genomics) pipeline. *Cellranger mkfastq* converted Illumina bcl files to fastq files then *Cellranger count* aligned the fastq files to mouse (mm10) or human (GRCh38)(GENCODE v32/Ensembl 98) reference genomes, filtered repetitive unique molecular identifiers (UMI) and generated a matrix containing gene counts in each cell. Following the Cell Ranger pipeline, the count matrices were imported into R and further analyzed using the software package Seurat v4(*34, 79*). Following the Seurat workflow, cells were filtered ensuring that contained a minimum of 200 genes and a maximum of 5000 genes, as well as ensuring that the genes present be expressed in at least 3 cells. Cells that expressed a high number of mitochondrial genes (greater than 10% of the total genes) were also removed. Following the sub-setting, the count data was log normalized (Seurat’s *NormalizeData* function) and scaled (Seurat’s *ScaleData* function) to correct for effects that highly expressed genes can have on the data set.

### Spatial Transcriptomics

The sentinel lymph node of human melanoma patients was biopsied, embedded in OCT, and frozen. 10 μm sections were generated using a cryostat and mounted on barcoded 10x Genomics slides. Barcoded cDNA libraries were generated using the Visium Spatial Gene Expression Slide & Reagent Kit (10x Genomics) at the Duke University Molecular Genomics Core. Tissues were stained with H&E to allow for mapping of spatial transcriptomics to the lymph node tissue. cDNA libraries were sequenced on an Illumina Novaseq 6000 sequencer using 50,000 read pairs per tissue covered spot on the capture area. The following read parameters were used: Read 1: 28 Cycles, i7 Index Read: 10 Cycles, i5 Index Read: 10 Cycles, Read 2: 90 Cycles. Samples were demultiplexed and aligned to the GRCh38 (GENCODE v32/Ensembl 98) human reference transcriptome using the Space Ranger 2.0 pipeline (10x Genomics). The Loupe Browser v5.0 (10x Genomics) was used to select capture spots that contained lymph node tissue. Further analysis was performed using Seurat v5. Data was normalized using Seurat’s *SCTransform* function and the top 30 principal components were used for PCA and UMAP dimensional reduction.

### scATAC-seq Processing and Analysis

For scATAC-seq experiments, 5,000 DCs were sorted from the tumor draining lymph node (inguinal) of BRAF^V600E^ PTEN^-/-^ melanoma bearing mice using FACS as previously described. Cell membranes were lysed and nuclei were isolated using a solution containing 0.1% Tween-20, 0.1% NP40 substitute, and 0.01% digitonin according to 10x genomics Nuclei isolation protocol. Single nuclei were encapsulated in gel bead-in-emulsions (GEMs) using the Chromium Next GEM Single Cell ATAC Library and Gel Bead Kit v1.1 (10x Genomics, Cat. PN-1000176) according to the manufacturer’s instructions. Quality control of libraries was performed using a Bioanalyzer and the final NGS libraries were quantified using NEBNext Library Quant Kit for Illumina (New England Biolabs, Cat. E7630L). For NGS, samples were sequenced on an Illumina NovaSeq 6000 at a depth of ∼25,000 reads per nucleus.

For data analysis, sequencing reads were processed using the Cell Ranger ATAC 2.0 pipeline (10x Genomics). Reads were first demultiplexed and fastq files were generated from raw Illumina bcl files using the command *cellranger-atac mkfastq.* Then, *cellranger-atac count* was used to align fastq files to the mouse (mm10) reference genome (GENCODE v32/Ensembl 98), filtered repetitive unique molecular identifiers (UMI) and generated a matrix containing accessible chromatin counts in each nuclei. The Seurat extension Signac was used for further quality control and processing of the scATACseq count-nuclei matrix. In brief, unique counts were required to occur in at least 10 nuclei and each nuclei required at least 200 features. Nuceli were then subset further to exclude low quality cells. For this, peaks were called and the total number of fragments in peaks was determined. Nuclei with fewer than 3000 and greater than 20,000 fragments were omitted to omit nuclei with poor sequencing depth and potential doublets or clumps of nuclei. Also, at least 15% of the total counts were required to fall within ATAC-seq peaks. Data was then normalized using TF-IDF and UMAP was used for dimensional reduction.

### Cellular Metabolism Analysis

For Seahorse XF Mito Stress Test assay, BMDCs were isolated and cultured from either CD11c-Cre-Srebf2^fl^ or wild type littermates and plated at a concentration of 1 x 10^5^ cells per well on a poly-d-lysine-coated 96 well Seahorse plate in Seahorse XF RPMI medium containing 1 mM pyruvate, 2 mM glutamine and 10 mM glucose. Prior to running the assay BMDCs were incubated in a non-CO2 incubator at 37° C for 1 hour. Throughout the assay, the oxygen consumption rates (OCR) and extracellular acidification rates (ECAR) of BMDCs were measured in response to 1.5 μM oligomycin, 1.0 μM FCCP and a combination of 0.5 μM rotenone/antimycin-a using a Seahorse XFe96 Extracellular Flux Analyzer (Agilent).

For the SCENITH assay, the tumor draining lymph nodes were isolated from BRAF^V600E^ PTEN^-/-^ melanoma-bearing mice. TDLNs were dissociated by incubation in RPMI containing collagenase and DNase for 30 minutes at 37° C. LN cells were washed twice and plated in RPMI containing 1 mM pyruvate, 2 mM glutamine and 10 mM glucose at 1 x 10^6^ cells/mL, 0.5mL/well in 48-well plates. The cells were treated with vehicle control or the metabolic inhibitors 2-Deoxy-D-Glucoae (2-DG, 100 mM), oligomycin (oligo, 1 μM) or etomoxir (eto, 40 μM) for 45 minutes. Puromycin (puro, 10 μg/mL) was added to the cells for the final 15 minutes of the assay. Cells were then washed twice in PBS and were proceeded to the flow cytometry staining protocol. Next, cells were stained in LIVE/DEAD Fixable Aqua Dead Cell Stain Kit (30 min room temperature 1:1000 dilution). Cells were washed 2x in FACS buffer (5% FBS, 2 mM EDTA in PBS), FC receptors were blocked using anti-CD16/CD32 (BD Biosciences, Cat. 553142) for 15 minutes at 4° C and surface proteins were stained to identify dendritic cell sub-populations for 1 hour at 4° C. Cells were then fixed and permeabilized using the eBioscience Foxp3/Transcription Factor Staining Buffer Set (ThermoFisher). Cells were then stained using an antibody that recognized incorporated puromycin (Kerafast, Inc) that has been labeled with AlexaFluor 594 Conjugation Kit (Abcam) for 1 hour. Cells were washed in FACS buffer and analyzed on a BD LSRFortessa (Duke Cancer Institute Flow Cytometry Core). For complete SCENITH Assay protocol(*45*).

### BODIPY Lipid Content Analysis

Neutral lipid content and lipid oxidation were probed using BODIPY 493/503 (ThermoFisher Scientific, Cat# D3922) or BODIPY 581/591 C11 (ThermoFisher Scientific, Cat# D3861), respectively. Following surface flow cytometry staining, cells were incubated with either BODIPY 493/503 or BODIPY 581/591 C11 at a concentration of 500 ng/ml in PBS for 30 minutes at 37° C. The cells were then washed 2x in PBS and resuspended in FACS buffer for flow cytometry analysis. BODIPY 493/503 was analyzed using a 488 nm excitation laser and a 525/50 emission filter. The reduced form of BODIPY 581/591 C11 is analyzed using a 561 nm emission laser and a 586/15 filter while the oxidized form is analyzed using a 488 nm excitation laser and a 525/50 emission filter.

### Treg Differentiation Assay

CD63^+^ DCs or CD63^-^ control DCs isolated from the LNs of Balb/c mice were sorted by FACS as described previously and cultured with naïve CD4^+^ T cells magnetically isolated from Fox3-GFP^+^ mice (C57Bl/6 background) using EasySep Mouse Naïve Cd4^+^ T cell isolation kit (Stemcell Technologies, Cat# 19765) at a ratio of 1:5 DC:T cell for 5 days in RPMI containing 10% FBS, 1% anti-anti, 2-mercaptoethanol, and 200 pg/ml TGFβ (R&D Systems, Cat# 7666-MB-005/CF). For IDO1 inhibition assays, CD63^+^ DCs or CD63^-^ control DCs were isolated as above and pre-treated with epacadostat (200 nM) for 8 hours prior to washing and co-culture with naïve CD4^+^ T cells for at a ratio of 1:5 DC:T cell for 5 days. Following the co-culture, the percentage of Foxp3-GFP^+^ Tregs was analyzed by flow cytometry.

### OT-I T Cell Proliferation Assay

The carboxyfluorescein succinimidyl ester (CFSE) dilutional assay was used to quantify T cell proliferation to determine the ability of CD63^+^ DCs to cross-present tumor-associated antigens, as well as to establish the extent to which CD63^+^ DCs interfere with antigen cross-presentation by other neighboring DCs. To test DC cross-presentation, CD63^+^ DCs or CD63^-^ control DCs isolated from the TDLNs of C57Bl/6 mice were sorted by FACS, as described previously, from mice bearing implanted BRAF^V600E^ PTEN^-/-^ melanomas that express ovalbumin. Naïve CD8+ T cells were isolated from OTI mice by magnetic selection (R&D Systems, Cat# MAGM207) labeled with cell trace CFSE (ThermoFisher Scientific, Cat# C34554) and co-cultured with the sorted DCs at a 5:1 ratio for 72 hours. Flow cytometry was then used to assess CFSE dilution and CD44 expression in CD8^+^ T cells.

To test if CD63^+^ DCs can interfere with the cross-presentation by other DCs, BMDCs were cultured and pulsed with ovalbumin (250 μg/ml, Millipore Sigma, Cat# A5503) overnight and washed 2x in RPMI with 10% FBS prior to co-culture. Naïve OTI CD8+ T cells were isolated and labeled with CFSE as above described. CD63^+^ or CD63^-^ DCs were sorted by FACS from the TDLN of BRAF^V600E^PTEN^-/-^ melanoma bearing mice. OVA-pulsed DCs were cultured with OTI T cells in a 1:5 ratio, respectively. CD63^+^ or CD63^-^ control DCs were added to the culture in ratios ranging from 1:10 to 1:40 relative to the OVA-pulsed DCs. The cells were incubated for 72 hours at which point the cells were analyzed by flow cytometry for CFSE dilution and expression of CD44 by CD8^+^ T cells.

### CRISPR-Cas9 Melanoma Cell Line Editing

Multiple gRNAs targeting *Slc16a1* (MCT1) were selected using the webtool CHOPCHOPv3 and cross-referencing to previous studies(*80*). gRNA sequences were cloned into a gRNA expression lentiviral vector (Addgene: #162076). To generate Cas9 expressing melanoma cells, BRAF^V600E^ PTEN^-/-^ melanoma cells were transduced with lentivirus containing the pSCAR-Cas9-blast-GFP vector (Addgene: #162074). This system, selective Cas9 antigen removal, (SCAR) allows for the removal of Cas9 antigen reducing the possibility of immune rejection due to Cas9 antigen(*81*). Following the generation of Cas9 expressing BRAF^V600E^ PTEN^-/-^ melanoma cells, gMCT1 lentivirus was generating by co-transfecting 293T cells in a 1:1:1 molar ratio of the gRNA expressing transfer vector and the packaging plasmids pSPAX2 and PMD2.G. Virus was collected filtered using a 0.45 micron filter and added to Cas9 expressing BRAF^V600E^ PTEN^-/-^ melanoma cells at a 1:1 ratio with fresh media. Three days following the transduction, cells were selected with blasticidin (5 μg/ml). To remove the Cas9 antigen, cells were transduced with an integration deficient lentivirus expressing Cre recombinase at an M.O.I. of 2.0 (Addgene: #162073). Cells with Cas9 removal were selected by sorting for GFP^-^ cells. Clonal cells were generated and MCT1 knockout as well as Cas9 Cre-mediated removal was verified by Western blot.

### CD45.1 DC Adoptive Transfer

cDC1s and cDC2s were enriched from the spleen of CD45.1 expressing mice on a C57Bl/6 background (B6.SJL-Ptprca Pepcb/BoyJ) using negative magnetic bead selection and sorted using FACS on a Sony SH800 Cell Sorter. 5 x 10^5^ cDC1s or cDC2s were injected intratumorally into BRAF^V600E^ PTEN^-/-^ transgenic melanomas that were approximately 800 mm^3^ in volume. 3 days following the intratumoral injection, tumors and lymph nodes were harvested and analyzed with flow cytometry for CD45.1 expressing DCs using the above described protocols.

### Mouse Tumor Studies

For inducible melanoma studies, melanoma was induced by injecting B6.Cg-Tg(Tyr-cre/ERT2)13Bos BRAF^tm1Mmcm^ Pten^tm1Hwu^/BosJ (BRAF^V600E^ PTEN^-/-^ Tyr::CreERT2, Strain #013590) mice subdermally with 4-hydroxytamoxifen (4-HT)(Millipore Sigma, H6278; 38.75 μg at the base of the tail. Once tumors reached approximately 75 mm^3^, mice were randomly assigned to a group. For fatostatin treatments, mice were treated twice weekly with Fatostatin (30 mg/kg, 150μl, 5% DMSO) via intraperitoneal injection.

For syngeneic tumor implant studies, 5 x 10^5^ BRAF^V600E^ PTEN^-/-^ melanoma cells were implanted at the base of the tail of either *Srebf2^loxp/loxp^* Itgax-Cre mice or littermates lacking Itgax driven Cre on a C57Bl/6 background. Tumor growth was measured using calipers twice weekly. The equation: volume^3^ = [(length)(width)^2^]/2 was used to calculate tumor volumes for all experiments.

### Statistical Analysis

Graphpad Prism 9 (Graphpad Software) was used for statistical analysis. An unpaired 2-tailed student’s *t* test was used to compare mean differences between two groups. Univariate 1-way ANOVA followed by Sidak’s post hoc multiple comparison’s test was used to compare mean differences between three or more groups. A *P* value of less than 0.05 was considered significant. Quantitative data was presented as mean ± SEM.

### Study Approvals

All mouse experiments were performed according to an IACUC-approved protocol at Duke University Medical Center. All patients were provided written informed consent under an IRB-approved protocol at Duke University Medical Center and the Duke Cancer Institute (Pro00090678).

## List of Supplementary Materials

Figs. S1 to S8

Table S1 and S2

Data File S1 and S2

GEO Files

## Author Contributions

M.P.P. and B.A.H. conceptualized the project, designed all experiments, and analyzed all data. M.P.P. either performed or directed all experiments. Y.X., Y.N., N.Y., X.Y., B.T., A.H. assisted with experiments and provided technical support. G.B. provided clinical resources for the project. B.A.H. supervised all experiments. M.P.P. and B.A.H. wrote the manuscript. B.A.H., B.T., N.Y., N.C.D., A.H., G.B. reviewed and edited the manuscript.

## Acknowledgements

The authors would like to thank the Duke Cancer Institute Flow Cytometry Shared Resource, the Duke Molecular Genomics Core, and the Duke Center for Genomic and Computational Biology. We thank Yongjuan Xia in the University of North Carolina Pathology Services Core (PSC) for expert technical assistance with Histopathology and Digital Pathology. The PSC is supported in part by an NCI Center Core Support Grant (P30CA016086). This work was supported in part by a NIH/NCI 5F32CA247067 (to M.P.P.), a Duke Cancer Institute Pilot Award, 5P30CA14236 (to B.A.H.), a DoD Idea Award, W81XWH2110847 (to B.A.H), a Duke University Health Scholar Award (to B.A.H.), a Duke Strong Start Award (to B.A.H.), a Damon Runyon Physician Scientist Award (to N.C.D.), and a NIH/NCI R37CA249085-02S1 (to B.T.).

## Supplementary Figure Legends

**Fig. S1. mregDCs express increased immunosuppressive and mevalonate pathway genes.** (**A**) Schematic showing the isolation of DCs by FACS from the LNs of tumor bearing or non-tumor bearing mice prior to scRNAseq analysis. (**B**) Gene expression overlaid on UMAP plots for the cDC markers, *Itgax, Flt3,*, *Zbtb46, Irf8, Irf4, Xcr1 and Sirpa* to identify conventional DC subsets. (**C**) Heatmap of scRNA-seq data demonstrating top-25 differentially expressed genes between cDC1s, cDC2s, and mregDCs (*n =* 2 independent experiments) (**D**) Significantly enriched hallmark gene sets expressed in mregDCs relative to cDC1s and cDC2s. (*n =* 2 independent experiments) (**E**) Average gene expression comparing scRNAseq replicates for all cells within the TDLN samples (*n =* 2 independent experiments).

**Fig. S2. CD63^+^ DCs are enriched in the LNs of tumor-bearing mice.** (**A**) Flow cytometry gating strategy to sort CD63^+^ DCs from LNs of tumor-bearing mice demonstrating CD63 gating using an APC isotype control. (**B**) Flow cytometry of DCs sorted from mice bearing Kras^G12D^ p53^-/-^ non-small cell lung carcinoma showing CD63^+^ DCs (*n =* 4). (**C**) Bar graph of the relative abundance of cDC1s, cDC2s, and CD63^+^ DCs in LNs spleen and tumor of BRAF^V600E^ PTEN^-/-^ melanoma-bearing mice (*n =* 4). (**D**) Scatter plot comparing the gene expression profiles of cDC1, cDC2 and mregDCs from melanoma TDLN to the gene expression profiles cDC1, cDC2 and mregDCs from tumor-free mice. (**E**) Flow cytometry gating strategy to identify CD45.1 DCs that have migrated from tumor to the TDLN (*n =* 4). Data is shown as mean ± SEM with individual data points. All data is representative of 2-3 independent experiments. * *P* < 0.05.

**Fig. S3. CD63^+^ mregDCs promote CD4^+^ T cell proliferation.** (**A**) Flow cytometry quantification of DQ-Ovalbumin uptake and processing by CD63^+^ mregDCs, cDC1s, and cDC2s (*n =* 3). (**B**) T cell proliferation assay of CFSE-labeled OTII T cells co-cultured with either CD63^-^ or CD63^+^ DCs isolated from the TDLN of mice bearing ovalbumin expressing melanomas (5:1 T cells: DC) (Top) Schematic of experiment (Bottom) Flow cytometry plots and quantification (*n =* 5). (**C**) Heatmap showing the differential expression of T_H_2-related genes in cDC subpopulations identified by scRNA-seq pathways (*n =* 2 independent experiments). (**A**) Statistical analysis performed by two-way ANOVA followed by Sidak’s multiple comparisons test. (**B**) Data analyzed using unpaired *t* tests. All data presented as a mean ± SEM. All data is representative of 2-3 independent experiments. * *P* < 0.05, \*\**P* < 0.005, \*\*\**P* < 0.0005.

**Fig. S4. Differential expression analysis of metabolic pathways in cDC subsets.** (**A** to **D**) Heatmaps generated from scRNA-seq data in **Fig. 1** showing the differential expression of genes associated with (**A**) the TCA Cycle, (**B**) Fatty acid metabolism, (**C**) Glycolysis, or (**D**) average expression across select metabolic pathways (*n =* 2 independent experiments).

**Fig. S5. Cellular metabolism analysis reveals the DC subpopulation dependencies on specific metabolic processes.** (**A** to **B**) SCENITH analysis of cDCs. (**A**) Representative flow cytometry histograms showing puromycin incorporation following treatment of cDCs with etomoxir, 2-deoxy-glucose, oligomycin, combination of the three inhibitors, or vehicle controls. (**B**) Normalized puromycin incorporation (gMFI) following treatment of cDC1s, cDC2s, and CD63^+^ mregDCs with etomoxir, 2-deoxy-glucose, oligomycin, combination of the three inhibitors, or vehicle controls. (**B**) Statistical analysis performed by two-way ANOVA followed by Sidak’s multiple comparisons test. All data presented as a mean ± SEM. \**P* < 0.05, \*\**P* < 0.005, \*\*\**P* < 0.0005.

**Fig. S6. Targeting SREBP2 in melanoma.** (**A**) MTS cell proliferation assay 24- and 48-hours following treatment of BRAF^V600E^ PTEN^-/-^ melanoma with fatostatin to determine the IC_50_ (*n* = 3). (**B**) qPCR analysis probing expression of SREBP2 target genes by DCs sorted from the LNs of melanoma-bearing mice following treatment with fatostatin (30mg/kg) or vehicle (*n* = 3). (**C**) Flow cytometry gating strategy to quantify CD4^+^FoxP3^+^ Tregs in the LNs of melanoma-bearing mice. (**D**) Flow cytometry gating strategy to quantify tumor-infiltrating CD8^+^ T cells in melanoma-bearing mice. (**B**) Data analyzed using unpaired *t* tests. All data presented as a mean ± SEM. \**P* < 0.05.

**Fig. S7. DC-specific knockout of SREBF2.** (**A**) Western blot showing expression of SREBP2 in BMDCs isolated from CD11c*-Cre*, *Srebf2^-/-^* mice compared to littermates lacking Cre expression (SREBF2^fl/fl^) (n = 2). (**B**) qPCR demonstrating the normalized gene expression of *Srebf2* in BMDCs isolated from CD11c*-Cre*, *Srebf2^-/-^* mice compared to littermates lacking Cre expression (SREBF2^fl/fl^) (*n = 3*). (**C**) SCENITH assay comparing puromycin incorporation in DCs from CD11c- *Cre*, *Srebf2^-/-^* mice and control DCs as a measure of ATP production (*n =* 8). (**D**) Flow cytometry analysis of BMDC neutral lipid content after isolation from CD11c*-Cre*, *Srebf2^-/-^* mice compared to littermates lacking Cre expression (SREBF2^fl/fl^) based on BODIPY 493/503 flow cytometry. (Right) Representative flow cytometry histogram of BODIPY 493/503 mean fluorescence intensity (MFI) (*n =* 4). (**E**) Western blot demonstrating CRISPR Cas9 editing of *Slc16a1* (MCT1) in BRAF^V600E^PTEN^-/-^ melanoma cells (n = 2). (**F**) Quantification of lactate release following CRISPR Cas9 editing of *Slc16a1* (MCT1) in BRAF^V600E^PTEN^-/-^ melanoma cells (*n =* 5). All data analyzed using unpaired *t* tests. All data presented as a mean ± SEM. \**P* < 0.05, \*\**P* < 0.005. \*\*\**P* < 0.0005.

**Fig. S8. Spatial transcriptomics of the sentinel lymph node isolated from a human melanoma patient.** (**A**) Expression scores for B cell, CD8^+^ T cell, CD4^+^ T cell, and CD4^+^ Treg overlaid on the Visium spatial transcriptomics spot map (*n =* 3 independent experiments). (**B**) Scatterplot showing the correlation of mregDC expression scores with B cell, CD8^+^ T cell, CD4^+^ T cell, and CD4^+^ Treg expression scores with associated Pearson correlation coefficients (*n =* 3 independent experiments).

