## Supplemental Figures for "A SREBF2-dependent gene program drives an immunotolerant dendritic cell population during cancer progression"

### Extended Data Figure 1

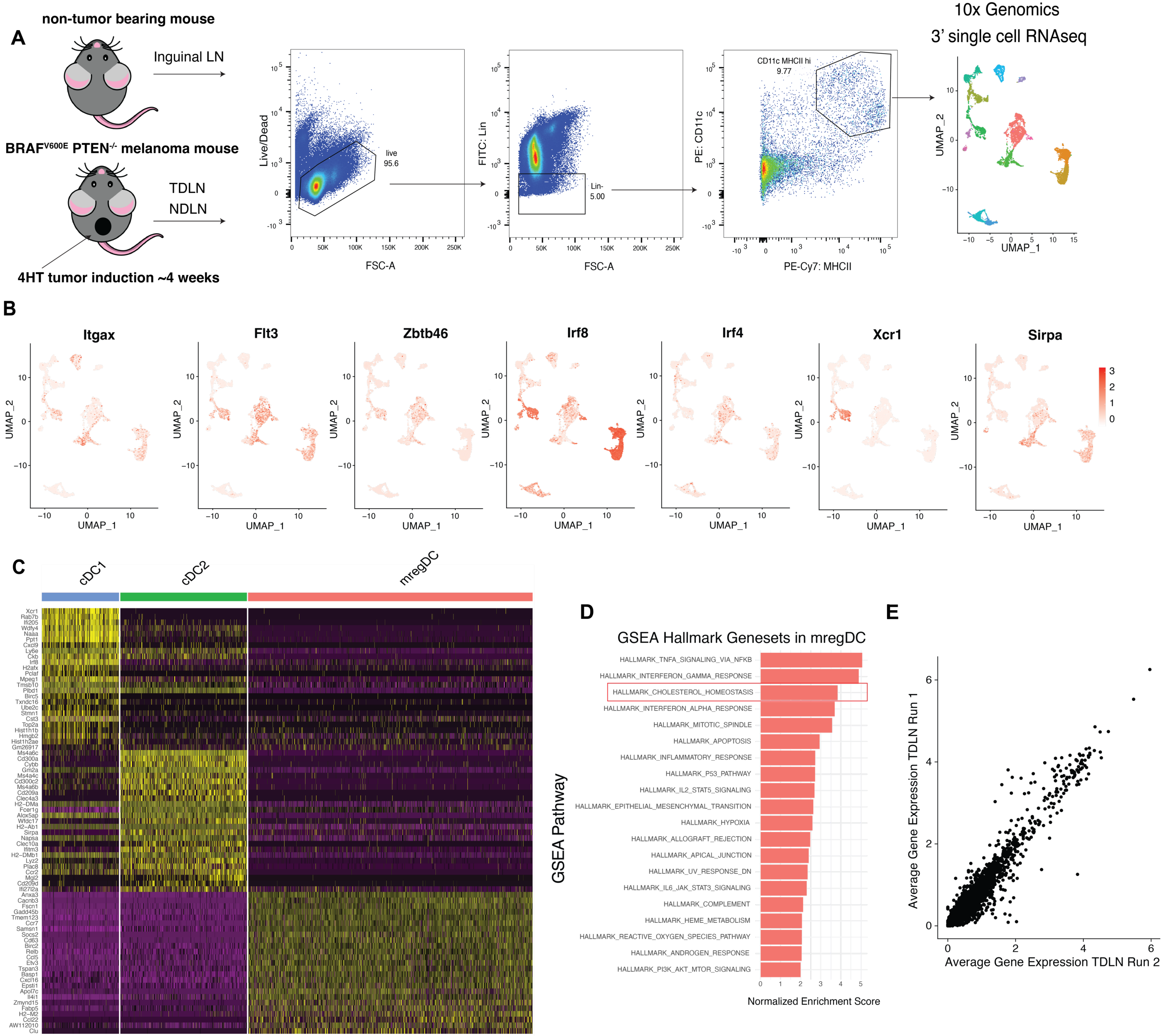

Extended Data Figure 2

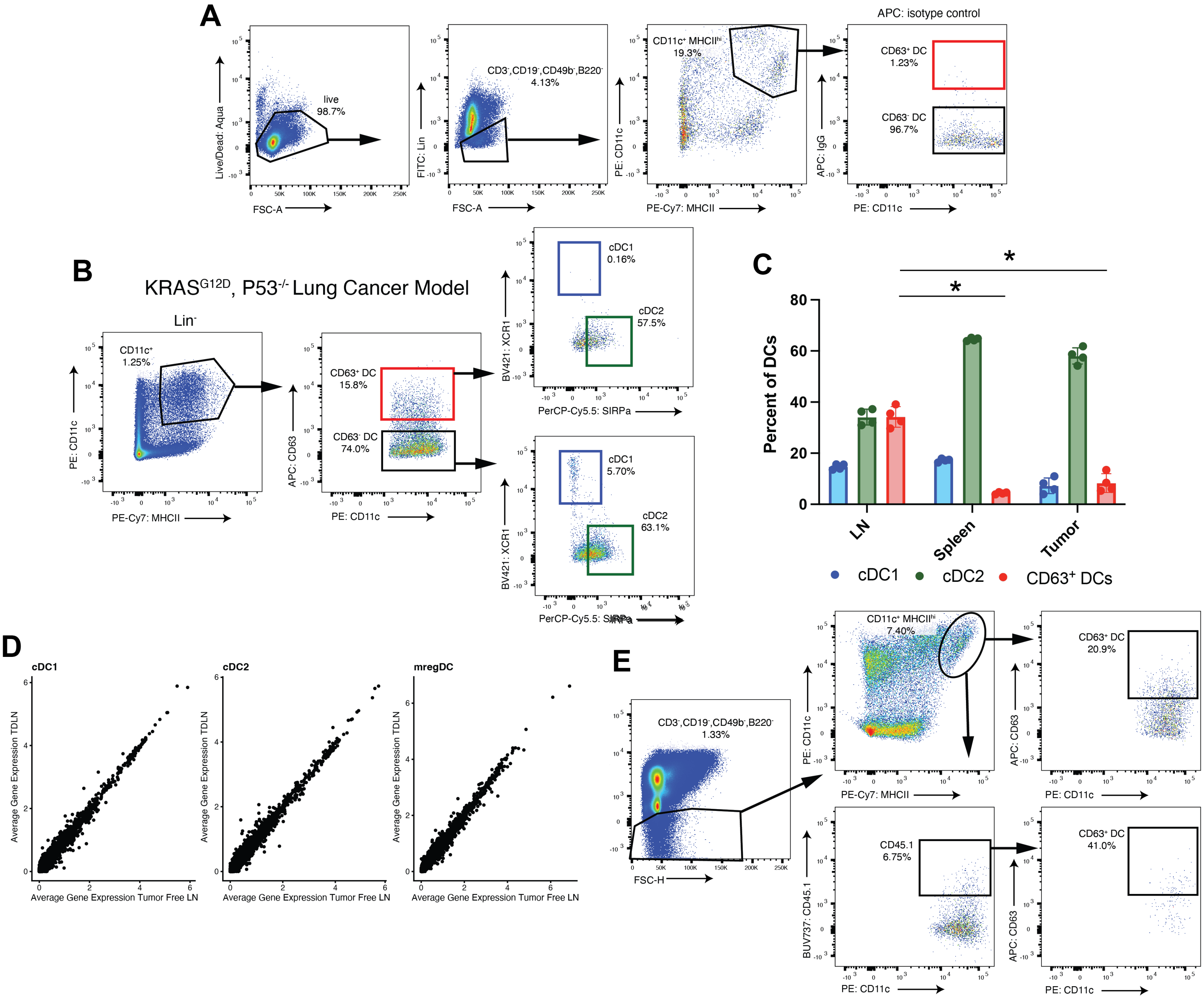

Extended Data Figure 3

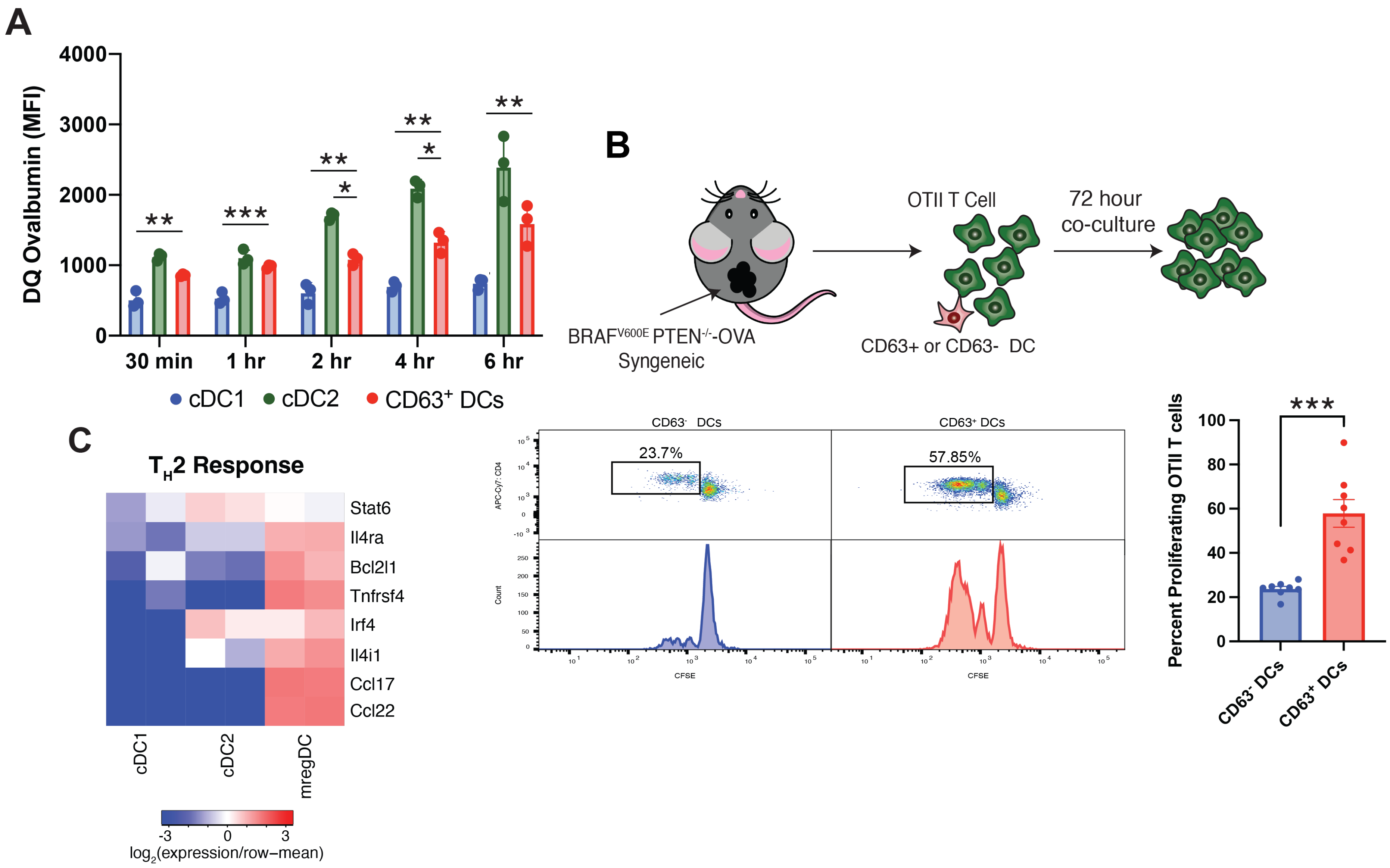

Extended Data Figure 4

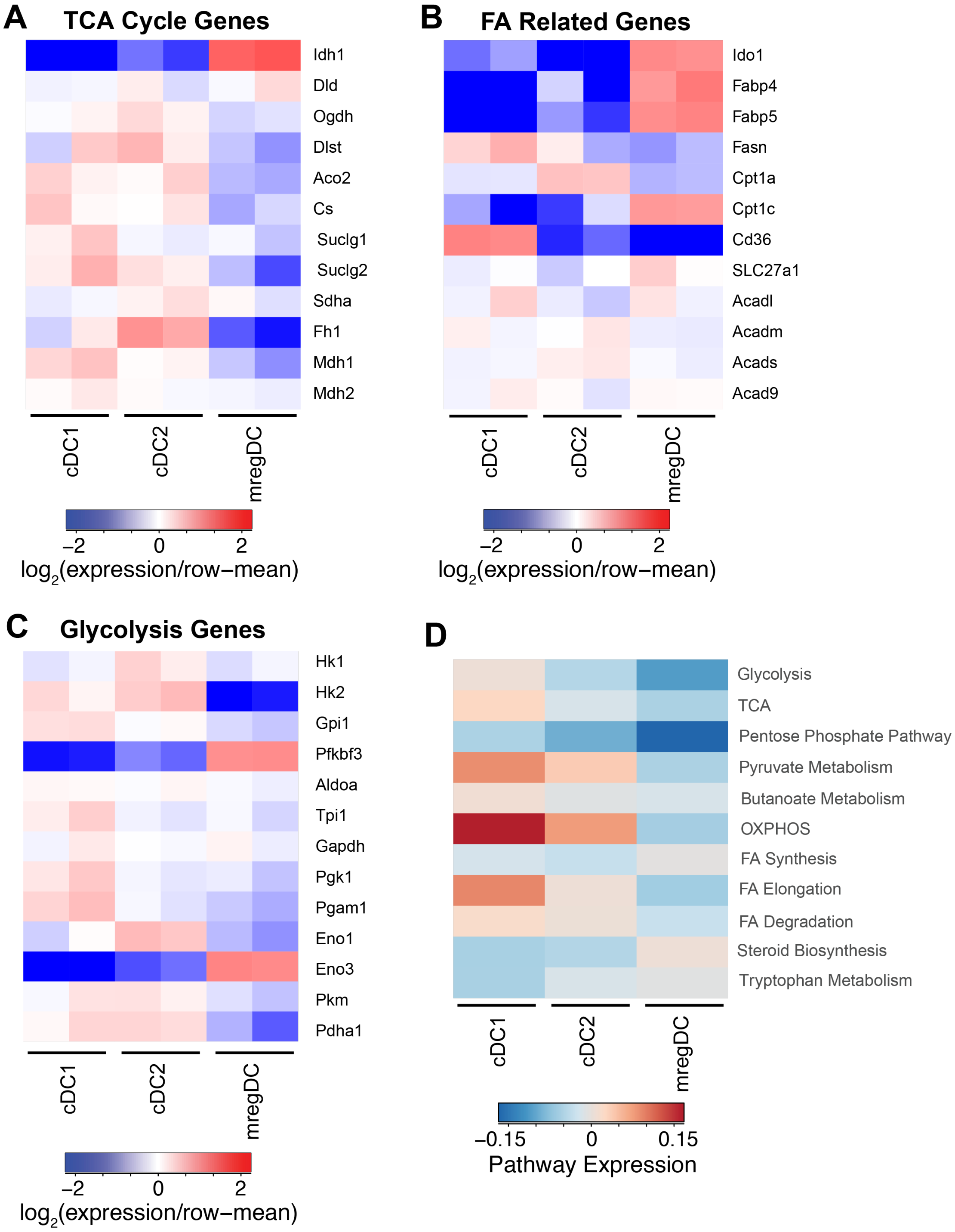

Extended Data Figure 5

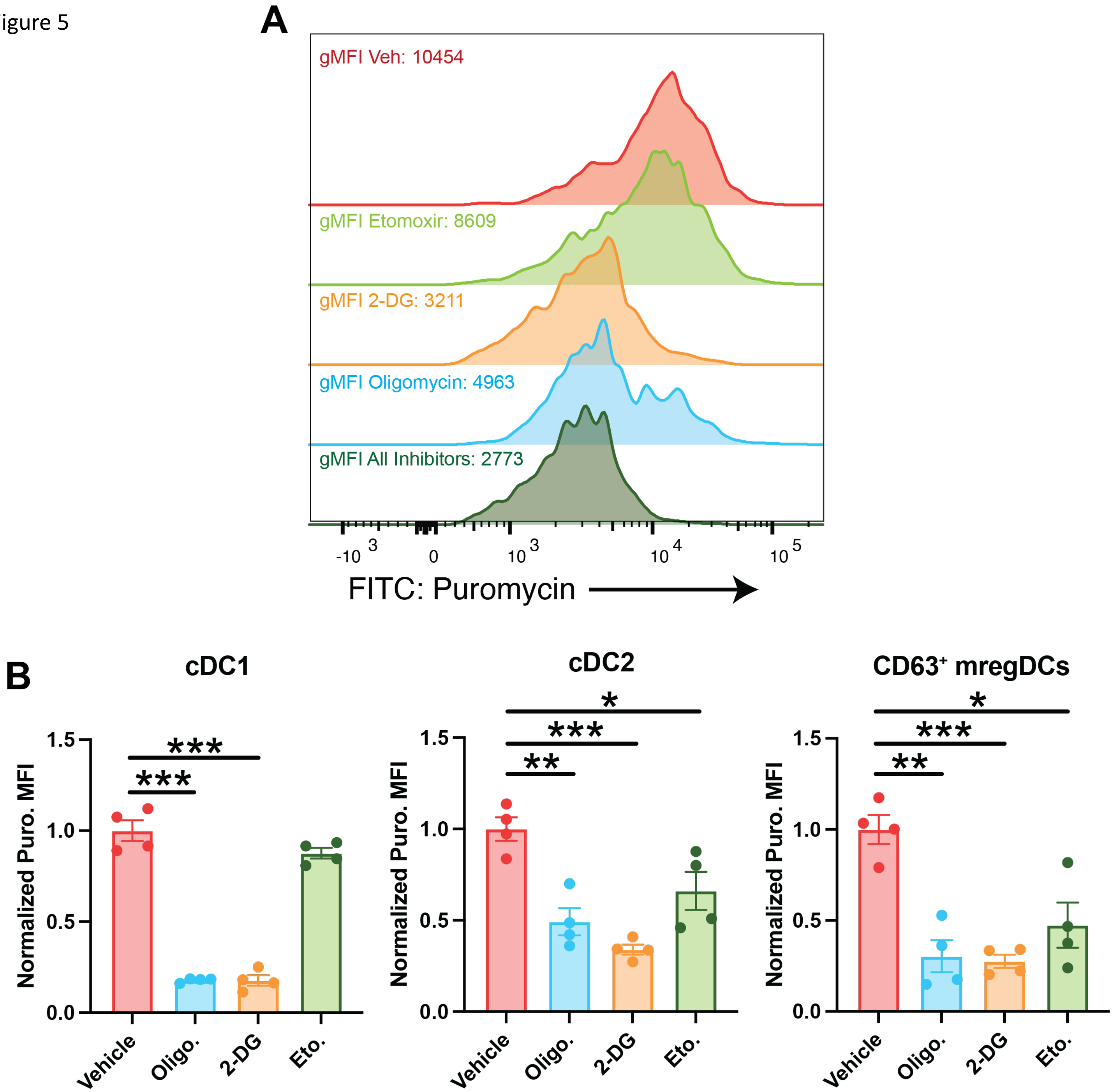

Extended Data Figure 6

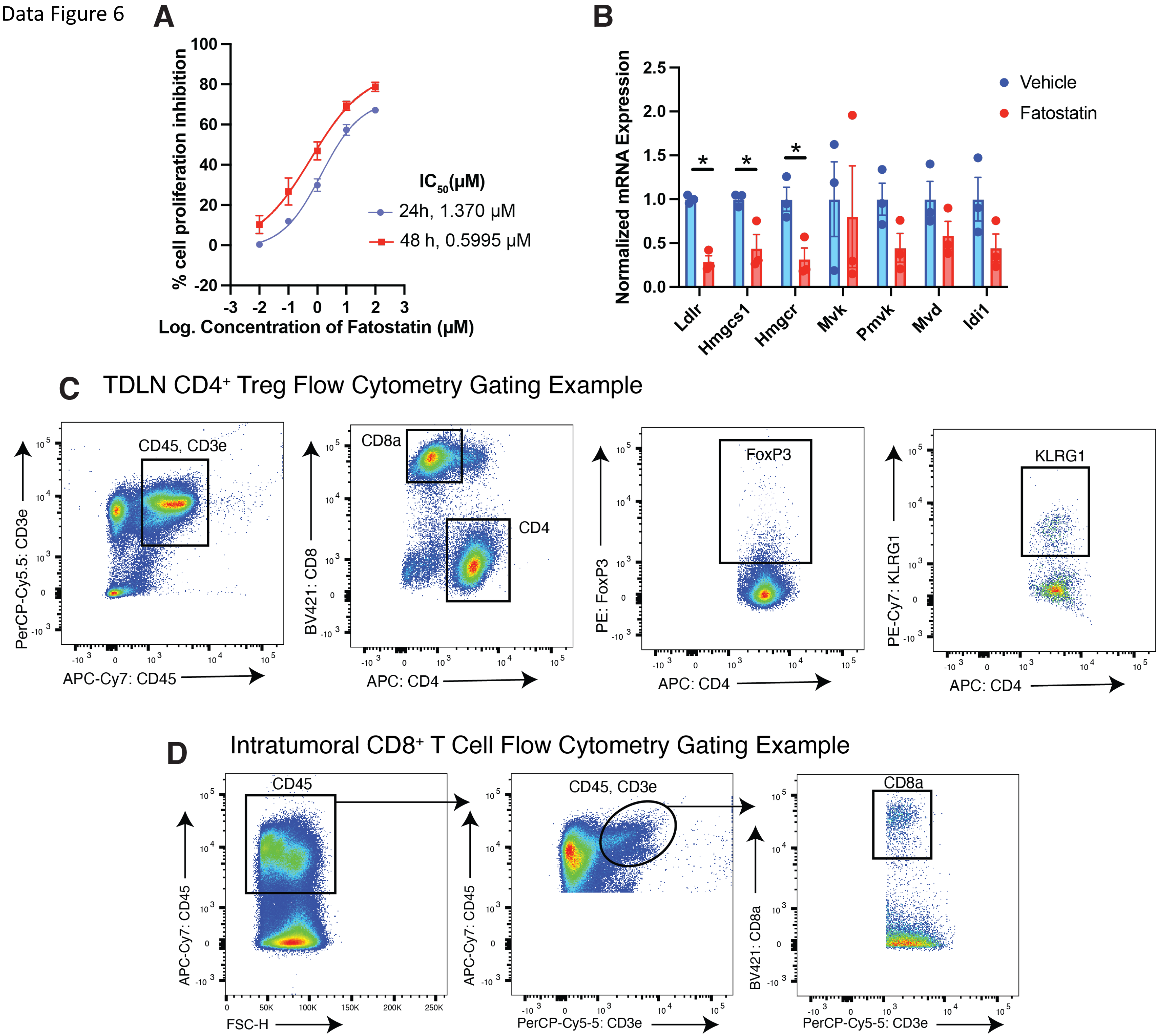

Extended Data Figure 7

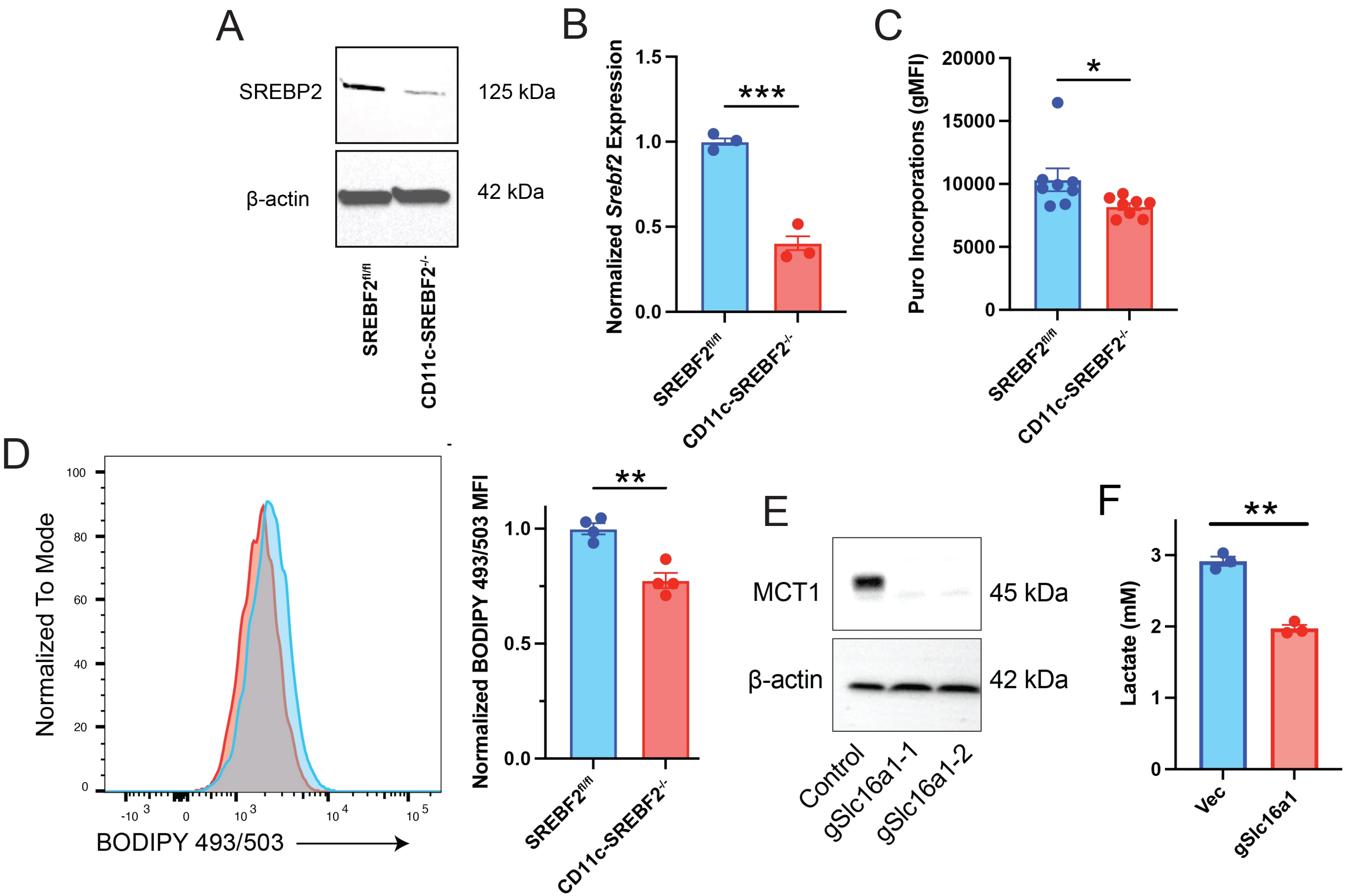

Extended Data Figure 8

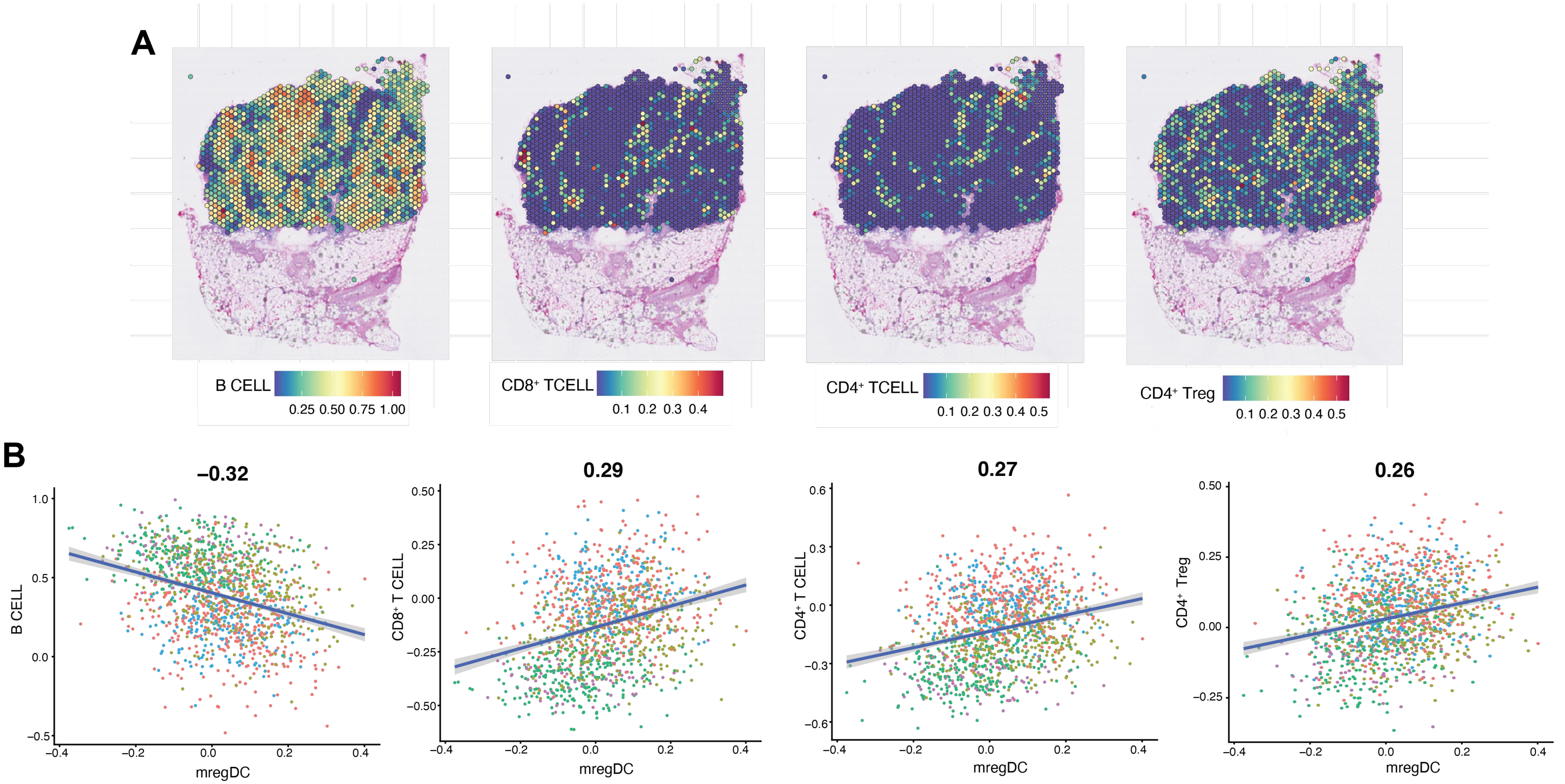
